# A *WFS1* variant disrupting acceptor splice site uncovers the impact of alternative splicing on β cell apoptosis in a patient with Wolfram syndrome

**DOI:** 10.1101/2023.12.29.573188

**Authors:** Raniero Chimienti, Silvia Torchio, Gabriel Siracusano, Valentina Zamarian, Laura Monaco, Marta Tiffany Lombardo, Silvia Pellegrini, Fabio Manenti, Federica Cuozzo, Greta Rossi, Paola Carrera, Valeria Sordi, Vania Broccoli, Riccardo Bonfanti, Giorgio Casari, Giulio Frontino, Lorenzo Piemonti

## Abstract

**Aims/hypothesis:** Wolfram Syndrome 1 (WS1) is an inherited condition mainly manifesting in childhood-onset diabetes mellitus and progressive optic nerve atrophy. The causative gene, WFS1, encodes for Wolframin, a master regulator of several cellular responses, whose mutations associate with clinical variability. Indeed, nonsense/frameshift variants correlate with more severe symptoms than missense/in-frame ones. As achieving a genotype-phenotype correlation is crucial to deal with disease outcome, works investigating the impact of transcriptional and translational landscapes stemming from such mutations are needed. Therefore, we sought to elucidate the molecular determinants behind the pathophysiological alterations in a WS1 patient carrying compound heterozygous mutations in WFS1 gene: c.316-1G>A, affecting the acceptor splice site (ASS) upstream exon 4, and c.757A>T, introducing a premature termination codon (PTC) in exon 7.

**Methods:** Bioinformatic analysis was carried out to infer the alternative splicing events occurring after disruption of ASS, followed by RNAseq and PCR to validate the transcriptional landscape. Patient-derived induced Pluripotent Stem Cells (iPSCs) were used as an in vitro model of WS1 and to investigate the WFS1 alternative splicing isoforms into pancreatic β cells. CRISPR/Cas9 technology was employed to correct ASS mutation and generate a syngeneic control for the ER-stress induction and immunotoxicity assays.

**Results:** We showed that patient-derived iPSCs retained the ability to differentiate into pancreatic β cells. We demonstrated that the allele carrying the ASS mutation c.316-1G>A originates two PTC-containing alternative splicing transcripts (c.316del and c.316-460del), and two ORF-conserving mRNAs (c.271-513del and c.316-456del) leading to N-terminally truncated polypeptides. By retaining the C-terminal domain, these isoforms sustained the endoplasmic reticulum (ER)-stress response in β cells. Otherwise, PTC-carrying transcripts were regulated by the nonsense mediated decay (NMD) in basal conditions. Exposure to cell stress inducers and pro-inflammatory cytokines affected the NMD-related gene SMG7 expression levels (>2 fold decrease; p<0.001) without eliciting a robust unfolded protein response in WFS1 β cells, thus resulting in a dramatic accumulation of the PTC-containing isoforms c.316del (>100-fold increase over basal; p<0.001) and c.316-460del (>20-fold increase over basal; p<0.001) and predisposing affected β cells to undergo apoptosis. Cas9-mediated recovery of ASS retrieved the canonical transcriptional landscape, rescuing the normal phenotype in patient-derived β cells.

**Conclusions/interpretation:** This study represents a new model to study Wolframin, highlighting how each single mutation of WFS1 gene can determine dramatically different functional outcomes. Our data point to increased vulnerability of WFS1 β cells to stress and inflammation, and we postulate that this is triggered by escaping NMD and accumulation of mutated transcripts and truncated proteins. These findings pave the way for further studies on the molecular basis of genotype-phenotype relationship in WS1, to uncover the key determinants that might be targeted to ameliorate the clinical outcome of patients affected by this rare disease.

## Introduction

Wolfram Syndrome 1 (WS1) is a progressive neurodegenerative disorder mainly characterized by co-morbidity of early-onset diabetes mellitus and optic nerve atrophy, in association or not with other endocrinological, urological and neurological abnormalities, with symptoms of highly variable onset, progression and severity (1, 2). Disease manifestations are attributable to pathogenetic variants of the *WFS1* gene, spanning 33.4kb of genomic DNA within the short arm of chromosome 4 (4p16). *WFS1* consists of eight exons, of which the first one is noncoding. The 3.6kb *WFS1* mRNA encodes for an 890 amino acids-long polypeptide, named Wolframin, having an apparent molecular weight of 100kDa. The protein displays subcellular localization to the endoplasmic reticulum (ER), where it plays a pivotal role in the control of the unfolded protein response (UPR), Ca^++^ homeostasis, ER-mitochondrial cross-talk and autophagy (3–6).

Wolframin structure and its interactome have not been entirely elucidated, but it has been reported to have protein-protein interaction domains at both N- and C-terminal, separated by nine transmembrane domains (TMDs) which allow the insertion in the ER membrane (7). Furthermore, the N-terminal domain is involved in the stabilization of tetrameric form, which is thought to be the most common and functional conformations of Wolframin (8). The 5’-untranslated region (UTR) of *WFS1* mRNA exhibits heterogeneity due to naturally occurring alternative splicing events (9, 10). More precisely, alternative splicing at the acceptor site of exon 2 causes a 4-bp deletion, enabling the differentiation of isoform 1 (aligned with the initial published sequence) from isoform 2, the latter starting four bases before the canonical ATG translation codon (9). Despite isoforms 1 and 2 generating the same protein, their biological significance remains unclear (11, 12).

To date, over 200 disease-causing mutations of *WFS1* have been identified along the entire sequence, with no evidence of mutational hotspots (https://www.ncbi.nlm.nih.gov/clinvar/?term=WFS1[gene], updated November 2023). As different *WFS1* variants mirror a wide range of clinical phenotypes, several genotype-phenotype association studies have been performed to aid clinicians in predicting more accurate prognoses and pave the way for personalized treatments (13–18). However, works investigating the molecular effects of specific *WFS1* variants on therapeutics are rare (19) and those proposing in-depth characterization of the transcriptional/translational outcomes stemming from the disruption of splice sites and/or introduction of premature termination codon (PTC) have not been performed yet.

It is well established that alternative splicing (AS) plays an important role in β cell function and viability (20, 21). Several alternative splicing variants contain PTC or intronic sequence after stop codon that can elicit nonsense mediated decay (NMD) to prevent the accumulation of aberrant transcripts or not-functional proteins (22). In pancreatic β cells, homeostatic pathways, including NMD and UPR, are involved in controlling and degradation of AS-deriving aberrant transcripts. Of note, several monogenic diabetes associated genes such as *HNF-1α* (23, 24), *GCK* (24), and *TCF7L2* (25) have AS variants, suggesting the importance of understanding the RNA surveillance mechanisms in pancreatic β cells to better explain disease phenotype (26).

In our work, taking advantage of both induced pluripotent stem cells (iPSCs) and CRISPR/Cas9 technologies, we sought to decipher the consequences at the transcriptional and translational level of two recently discovered pathogenic *WFS1* mutations: c.316-1G>A, affecting the acceptor splice site (ASS) upstream exon 4, and c.757A>T, introducing a PTC in exon 7 (27, 28). Finally, we explored the maturation of *WFS1* mRNA in patient-derived iPSCs and after their differentiation into pancreatic β cells, evaluating the impact of *WFS1* PTC-carrying transcript accumulation upon exogenous stress and inflammation-dependent NMD inhibition on survival of the insulin-producing β cells.

### Methods and Materials

#### PBMC isolation, reprogramming and cell culture

The healthy donor and patient-derived peripheral blood mononuclear cells (PBMCs) were obtained following informed consent and collected using Ficoll-Paque separation method on whole blood. WFS1 iPSCs were generated by reprogramming the CD34^+^ fraction from the patient-derived PBMCs using CytoTune-iPS 2.0 Sendai Reprogramming Kit (ThermoFisher) according to the manufacturer’s indications. The iPSC lines were routinely tested for mycoplasma contamination by using MycoAlert™ Mycoplasma Detection Kit by Lonza, according to the manufacturer’s instructions. Karyotyping was performed by ISENET Biobanking service unit in Milan, Italy. All WFS1-derived iPSC clones and their gene edited counterpart used in the present study had normal karyotype. Control iPSCs with WT genotype (CGTRCiB10) were already available and routinely employed in the laboratory. This WT iPSC line was used as unrelated healthy control and its in-depth characterization is reported in our previous publications (29, 30). Differentiation into pancreatic β cells was performed in adhesion following in vitro protocol published by our group (31) on subclones of WT iPSCs (#05, #B2, #T3), two clones of WFS1 iPSCs (#42 and #43) and three clones of WFS1^wt/757A>T^ iPSCs (#B10, #C6 and #F4). At the end of differentiation (day 24), iPSC-derived β cells were detached using 0.5 mM EDTA and displaced in 35 mm or 60 mm petri dishes on an orbital shaker at 55 rpm to aggregate into clusters (pseudo-islets). The pseudo-islets were maintained in suspension culture at 55 rpm until their use, using the following medium: CMRL 1066 (Mediatech) supplemented with 10% fetal bovine serum (FBS) (Lonza), 1% Penicillin/Streptomycin (Pen/Strep), 1% L-Glutamine (Euroclone), 10 μM Alk5i II, 1 μM T3, 10 mM Nicotinamide (Sigma), 10 μM H1152 (Euroclone), 100 U/ml DNase I (Sigma). Skin fibroblasts from a WS1 patient with genotype W189X/W189X and unrelated healthy control were provided by Dr. Vania Broccoli’s lab at IRCCS San Raffaele Hospital. Fibroblasts were kept in the Dulbecco’s modified Eagle’s medium (DMEM) with high glucose, supplemented with 10% FBS, 1% Pen/Strep, 1% L-Glutamine.

#### RNA extraction, retrotranscription and PCR/RT-qPCR

Total RNA was extracted by using mirVana Isolation Kit (Ambion) and quantified by BioTek Epoch Microplate Spectrophotometer (Agilent). About 1 µg of TURBO^TM^ DNAse (Thermo Fisher)-treated RNA was reverse transcribed using SuperScript™ IV First-Strand Synthesis System (Thermo Fisher), with oligo(dT) primers, according to the manufacturer’s instructions. For the identification of *WFS1* alternative splicing isoforms, 10 ng of cDNA was pre-amplified using *WFS1* primers listed in **ESM Table 1** (0.4 μM fw/rev) and the 1.25 U DreamTaq HS DNA Polymerase (Thermo Fisher), with 3.5 mM MgCl_2_ and 0.2 mM dNTPs, in 50 uL final volume reaction. Pre-amplification consisted of following cycling conditions: 1 x 95°C 3’, 15 x (95°C 30’’, 60°C 30’’, 72°C 45’’), 1 x 72°C 5’. A total of 1 uL of pre-amp reaction was used as template for further amplification. Second PCR was performed using 1.25 U DreamTaq HS DNA Polymerase, the same primers of pre-amp PCR (0.4 μM each), 2.0 mM MgCl_2_ and 0.2 mM dNTPs, with the following cycling conditions: 1 x 95°C 3’, 35 x (95°C 30’’, 60°C 30’’, 72°C 45’’), 1 x 72°C 5’. The PCR products were run on analytical pre-cast E-Gel™ Agarose Gels with SYBR™ Safe DNA Gel Stain, 2% (ThermoFisher), using the GeneRuler 100 bp Plus DNA Ladder (Thermo Fisher) as size markers. The spliced/unspliced (s/u) XBP-1 ratio

**Table 1.**
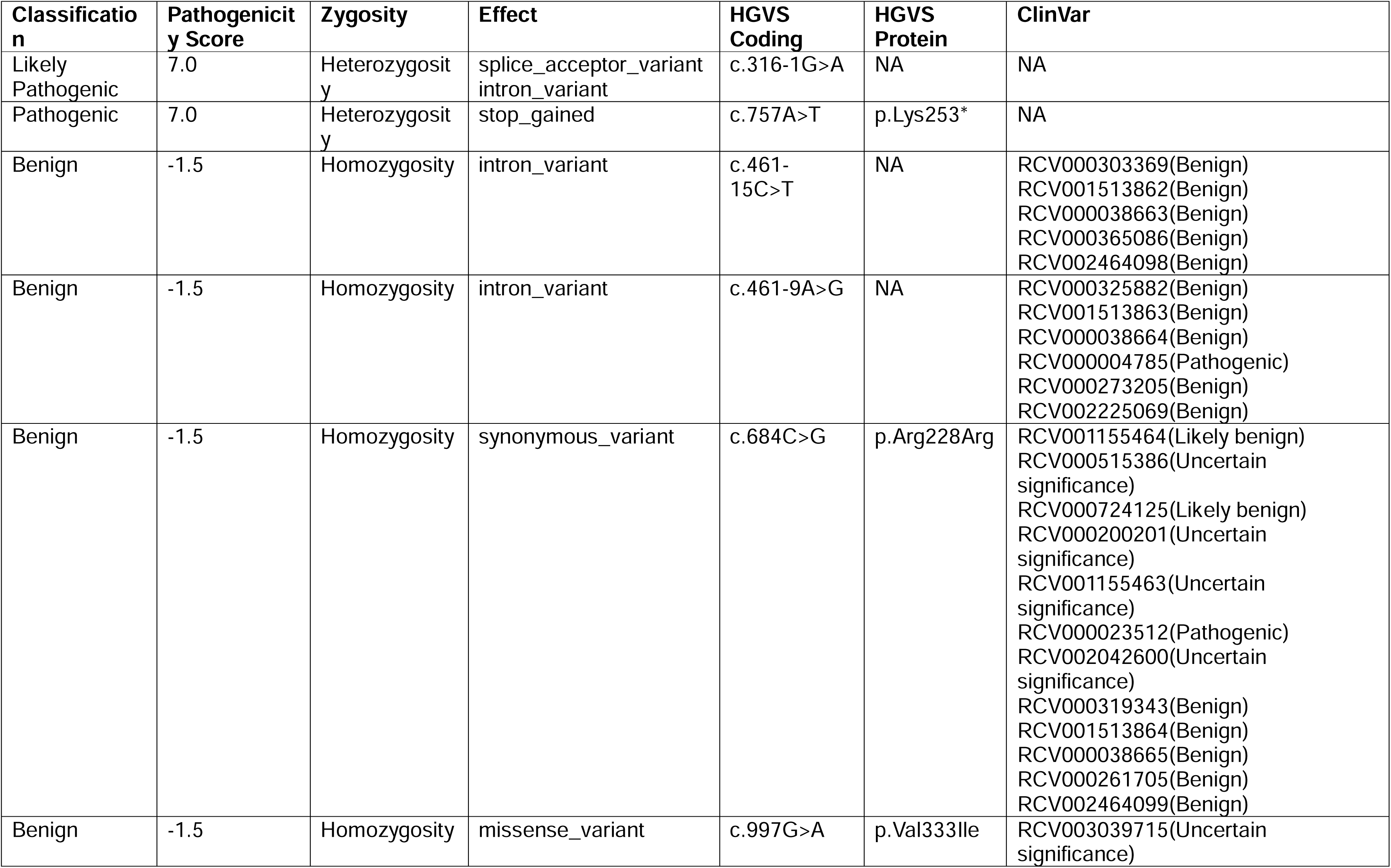
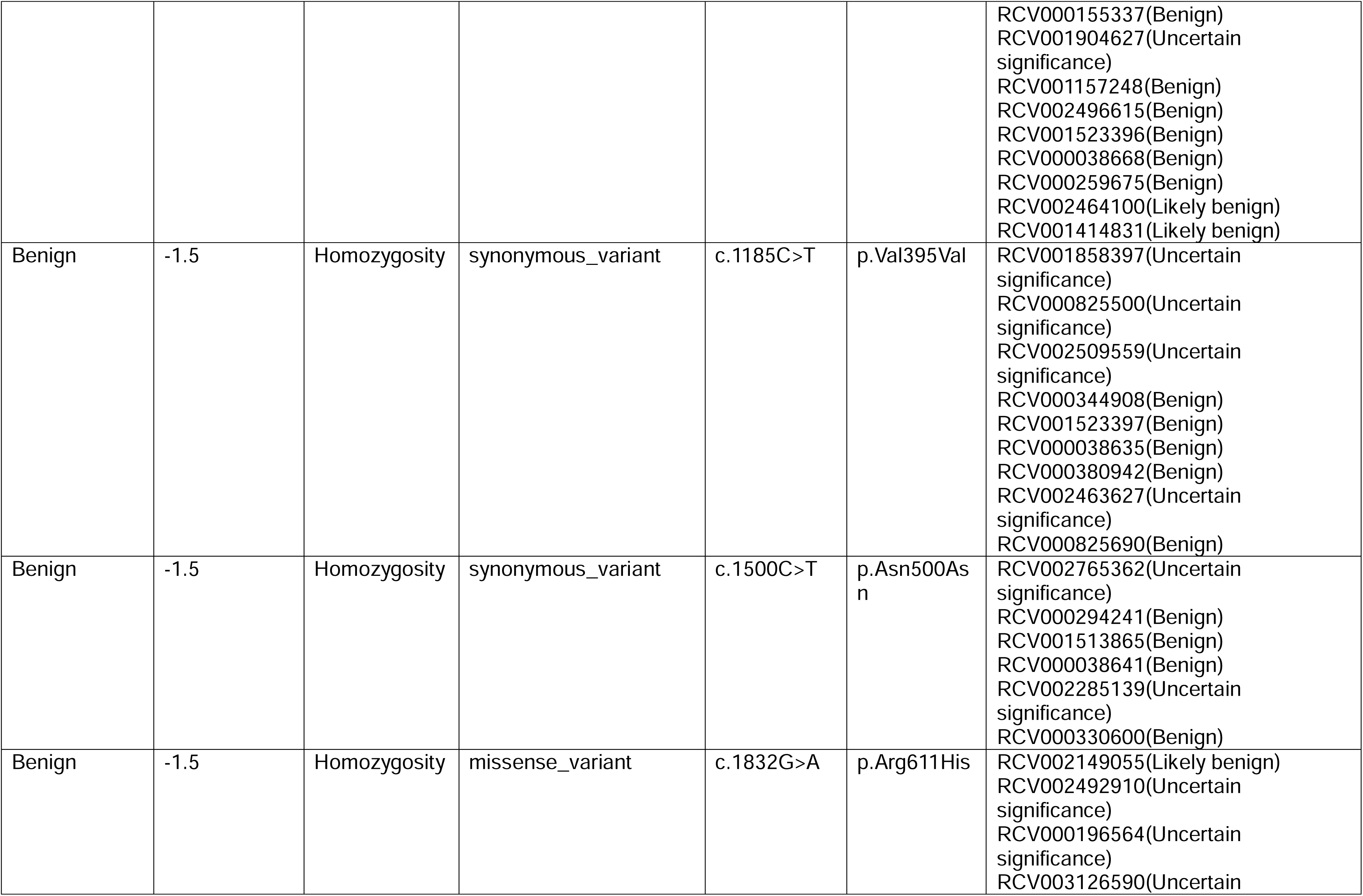
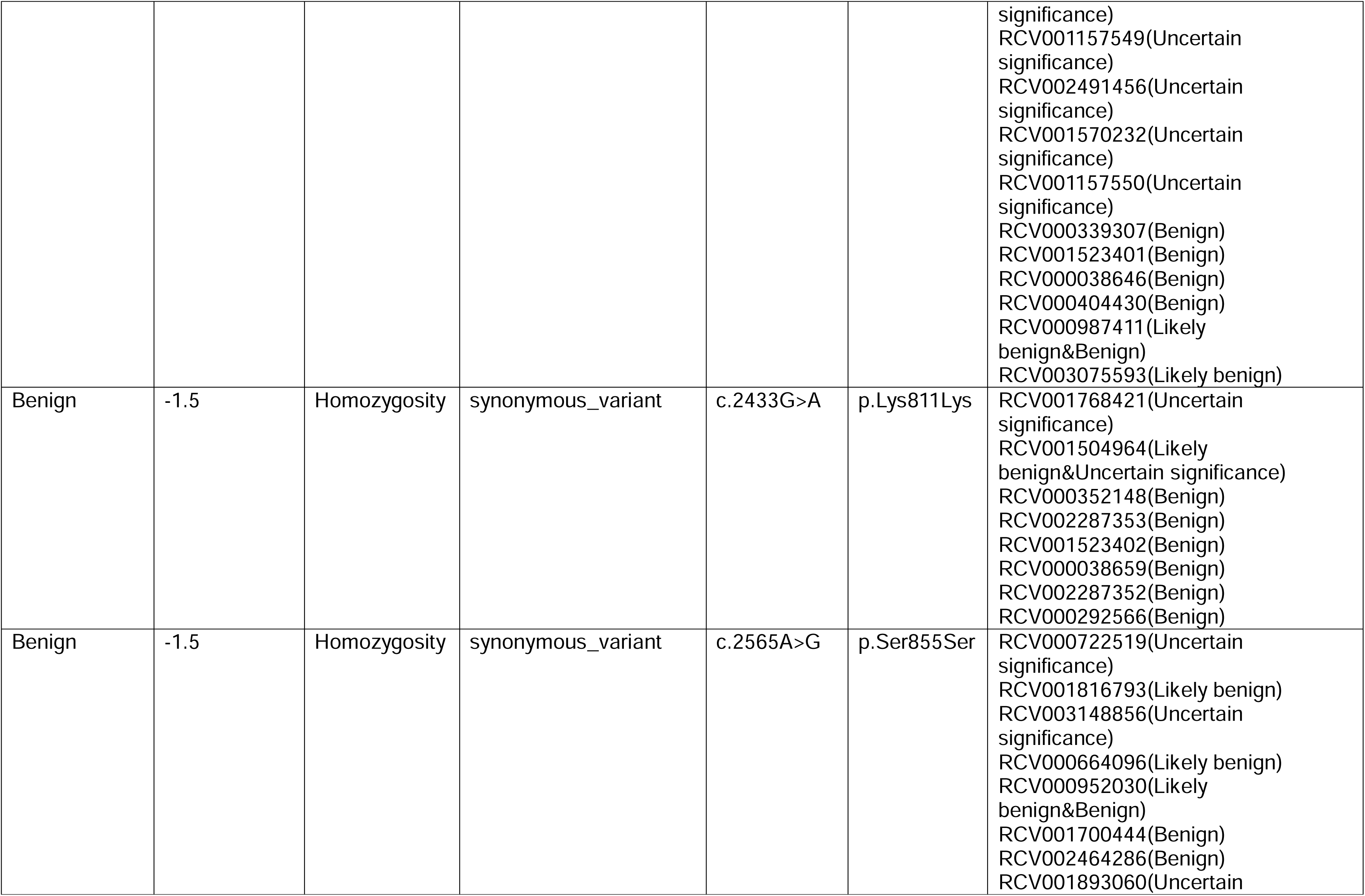
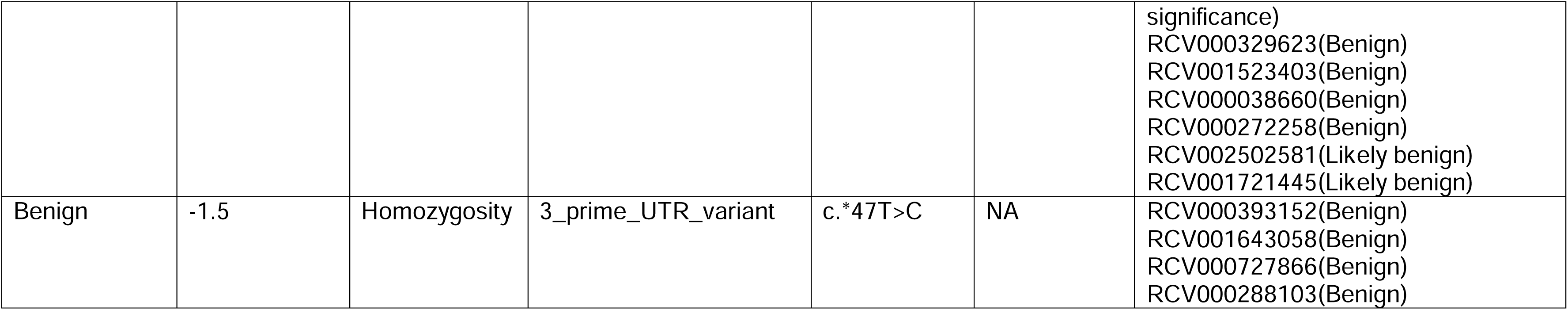
List of the WFS1 variants of the Wolfram syndrome patient as found by Next Generation Sequencing.

was measured by agarose gel electrophoresis and densitometric analysis. For RT-qPCR experiments, PowerUp Green Master Mix (Applied Biosistems) and TaqMan Universal PCR Master Mix (Applied Biosistems) were used according to the manufacturer’s protocol, and amplification with annealing temperature of 60°C was performed on a 7900HT Fast Real-Time PCR System (Applied Biosystem) with Fast 96-Well Block Module. Expression level of analyzed genes were normalized applying the 2^-ΔCt^ method, using GAPDH as housekeeping. A complete list of primers and TaqMan assay are reported in **ESM Table 1** and **ESM Table 2**.

#### PCR product purification, cloning and Sanger sequencing

WFS1-derived PCR products obtained following amplification with Ex3fw-Ex5rev primer pair were separated on 2% agarose extraction gel prepared dissolving low-melt-temperature agarose in 1X TBE. Both 324bp- and 180bp-long amplicons were purified separately by NucleoSpin PCR & Gel Clean-up (Macherey Nagel) and cloned into pCR4-TOPO TA vector using TOPO™ TA Cloning™ Kits for Sequencing (Thermo Fisher), following manufacturer’s instructions. The pCR4-TOPO construct was transformed into One Shot TOP10 (Thermo Fisher) chemically competent cells, then using LB-agar plates containing 100 µg/mL ampicillin as selection medium. The bacterial colonies were picked and analyzed for presence of insert by following amplification with the T3/T7 primers, and agarose gel electrophoresis. The pCR4-TOPO plasmid containing insert was purified using NucleoSpin Plasmid, Mini kit for plasmid DNA (Macherey Nagel). The Sanger sequencing was performed by the EZ-Seq service at Macrogen Europe. Electropherograms were visualized and sequence analyzed by using SnapGene v4.3.2 (Dotmatics).

#### Next Generation Sequencing

Following the patient’s referral to a geneticist for further examination, Next Generation Sequencing (NGS) was performed on a panel of 31 monogenic diabetes-related genes, including WFS1 (NM_006005). Enrichment of fragments was performed with the TruSight One Sequencing kit (Illumina). Sequencing of coding regions and exon-intron junctions was done on NextSeq Illumina platform. Diagnosis of Wolfram Syndrome 1 was confirmed when mutations in *WFS1* gene were reported in the medical record of the patients, after comparison with variants deposited in the following databases: NCBI dbSNP, 1000 Genomes, dbNSFP, ClinVar, LOVD. The deep sequencing analysis on PCR products obtained by amplifying *WFS1* sequence comprised between exon 3 and exon 5 was performed to confirm the sequence of the alternative splice variants deriving from c.316-1G>A-carrying allele in the WFS1 cells. PCR-derived fragments were purified by using NucleoSpin PCR & Gel Clean-up (Macherey Nagel). Libraries were obtained following the TruSeq Nano WGS (Illumina) protocol, deriving 250nt-long 4x10^6^ fragments per sample. Paired-end sequencing was performed through MiSeq_500_v2 Illumina platform on two WFS1 iPSCs clones (#01 and #42) and one WT iPSCs clone (#05). The reads in R1 and R2 FASTQ files generated at the end of the sequencing were analyzed for the base quality. Adapters were removed by using Trimmomatic, v.0.32. Preliminary, explorative alignment was performed by STAR v2.5.3a, mapping reads to the reference human genome hg38, Gencode version 28. Percentages of uniquely mapped reads were about 70-77% for the two WFS1 samples and 85% for the WT sample. More than 99.99% of reads were assigned to the *WFS1* gene for all the three samples.

#### CRISPR/Cas9-mediated gene editing

CRISPR/Cas9-mediated correction of the point mutation located upstream exon 4 of *WFS1* gene was performed in patient-derived iPSCs. GeneArt™ Precision gRNA Synthesis Kit (ThermoFisher) was employed for *in vitro* synthesis of the gRNAs, whose sequences are reported in **ESM Table 1**. As donor templates a single strand oligodeoxynucleotide (ssODN) homologous to the region encompassing the *WFS1* intron 3 - exon 4 boundary junction was used. 20 µg of Cas9 protein (TrueCut™ Cas9 Protein v2, ThermoFisher) and 4 µg of *in vitro* transcribed gRNAs were pre-incubated for 20 minutes at RT to form ribonucleoprotein (RNP) complex. An RNP:ssODN ratio of 1:1.5 was used by adding 36 µg of ssODN. Finally, 2 µg of pmaxGFP plasmid (Lonza) was added to the mix as a selection marker for cell sorting. For Cas9-mediated gene editing, 1·10^5^ iPSCs were electroporated by using the 4D-nucleofector with P3 electroporation solution (Lonza) and setting the CB-150 program, according to the manufacturer’s instructions. The electroporated cells were replated and cultured in complete fresh media for 5 days. Single cell cloning was performed following cell sorting of GFP-positive cells.

#### Cell sorting and FACS analysis

Electroporated cells were analyzed with FACSAria™ Fusion Flow Cytometer (BD) and GFP-positive cells were sorted in Vitronectin-coated 96-well plate for single cell cloning. Sorted cells were cultured in Essential 8 Flex supplemented with CloneR™ (Stemcell Tech.) until colonies emerged. Single clone-derived colonies were screened by restriction enzyme as described below. For intracellular staining, cells were fixed by using Cytofix/Cytoperm (Becton Dickinson, BD), then permeabilized with Phosflow™ Perm Buffer III (BD), according to the manufacturer’s instructions. Staining with conjugated antibody was performed for 30’ at 4°C. The list of antibodies used for FACS analysis is reported in **ESM Table 3**. For the ER-stress induction and immunotoxicity assay, early and late apoptosis were evaluated by using the FITC Annexin V Apoptosis Detection Kit I (BD), following the manufacturer’s instructions. Cells were inspected on CytoFLEX LX (Beckman Coulter) cytometer, and analyzed by FlowJo™ Software V.10 (FlowJo LLC, Ashland, Oregon, USA).

#### Homologous directed repair screening

Single colonies of growing clones were picked and transferred into direct PCR lysis buffer containing 150 mM NaCl, 10 mM Tris-HCl pH7.5, 5% Tween-20 and 0.4 mg/ml Proteinase K (Promega), then incubated for 6 hours at 55°C until complete lysis is achieved. Crude lysate was incubated for 45’ at 85°C to inactivate Proteinase K. After centrifugation to collect debris, 1 μL of lysate was used per 50 μL genotyping PCR reaction with HR fw/rev primers reported in **ESM Table 1**. The PCR product was purified using NucleoSpin PCR & Gel Clean-up (Macherey Nagel) and quantified by BioTek Epoch Microplate Spectrophotometer (Agilent). Up to 1 μg of DNA was digested with 10 U of SacI enzyme (NEB) in NEBuffer r1.1 for 1 hour at 37°C. Digested DNA derived from each clone was resolved on 2% agarose gel. Amplified sequence is 272 bp long. Enzymatic digestion of recombined sequence produces two bands of 152 bp and 119 bp. If homologous recombination with donor DNA involves one allele, the 272 bp-long and the 152/119 bp-long pair amplicons are observed (50% chance that the c.316-1G>A-carrying allele has been corrected). If homologous recombination occurs in both *WFS1* alleles, only digested products are displayed. For following experiments, only clones incurred in bi-allelic recombination were selected and evaluated for *WFS1* expression.

#### Immunofluorescence

For indirect immunofluorescence, cells were cultured on coated Falcon™ Chambered Cell Culture Slides. Cell monolayers were washed with PBS, fixed with 4% paraformaldehyde in PBS for 20’ at RT and then treated with 15 mM glycine for 5’ at RT. After blocking and permeabilization with PBS supplemented with 0.4% Triton X-100, 2% BSA, 5% FBS for 45’ at RT, cells were washed twice with PBS and then incubated overnight at 4°C with appropriate primary antibody reported in **ESM Table 3** diluted in PBS supplemented with 2% BSA. Cells were incubated with appropriate secondary antibody for 1h at RT. Nuclei were counterstained with Hoechst 33342. Images were acquired using Olympus FluoVIEW FV 3000 (VivaScope Research) confocal microscope and analyzed using the Fiji software v.1.52p (32).

#### Immunoblot analysis

Samples were lysed in M-PER™ and proteins were quantified by Pierce™ Rapid Gold BCA Protein Assay Kit. Protein extracts were resolved on SDS-PAGE (4-20%) using Novex™ Tris-Glycine Mini Protein Gels and electro-transferred onto a polyvinylidene difluoride (PVDF) membrane. Transfer efficiency was evaluated using Ponceau S (Sigma). After 1h of incubation in blocking buffer (TBS, 0.1% Tween 20, 5% skim milk) at RT, the membranes were probed overnight at 4°C with the primary antibodies reported in **ESM Table 3** diluted in TBS supplemented with 0.1% Tween 20 and 5% skim milk or BSA. Horseradish peroxidase-labeled antibodies (diluted 1:1000 in the same dilution buffer as the primary antibody) were used as secondary antibody, and revelation was performed using the SuperSignal™ West Pico PLUS chemiluminescent detection system according to the manufacturer’s instructions by ChemiDoc MP (Biorad). Quantification of protein levels was performed using Fiji software (32). GADPH was used as normalizer for protein quantification. All the listed reagents were purchased from ThermoFisher unless otherwise specified. Frozen EndoC-βH1 cell pellets were available in the lab (29) and were employed as controls of protein expression for β cells: they were resuspended in lysis buffer and subsequently processed as the other samples.

#### Non-sense mediated decay inhibition

To determine whether NMD inhibition can increase the PTC-carrying *WFS1* transcript levels, iPSCs were cultured in 6-well dishes and once reached ∼70% confluence were treated with 5 μM NMDI-14 (MedChemExpress) or with vehicle (DMSO 0.1%) for 16h. RNA was collected at the end of the treatment and processed following aforesaid methods.

#### ER-stress induction and immunotoxicity assay

For the thapsigargin (TG, Sigma)-induced ER-stress experiments iPSCs were treated for 2, 4, 8, 16 or 24h with 100 nM TG; for the cytotoxicity assay they were treated for 8h followed by 24 or 48h of recovery in fresh medium without TG. For iPSC-derived β cells, TG treatment lasted for a total of 8 or 16h with 50 nM TG before further analysis. The conditioning with inflammatory cytokines was performed up to 48h on iPSC-derived β cells by adding 50 U/mL rhIL-1β, 1000 U/mL rhIFN-ɣ and 10 ng/mL rhTNF-α (PeproTech) to the culture medium. Cells were analyzed by FACS for positivity to Annexin V/P.I. at 8, 16 or 48h, according to the protocol described above.

#### Hormone secretion

Ghrelin, glucagon and c-peptide levels were measured from culture supernatants by Bio-Plex Pro human diabetes kit (BioRad), following the manufacturer’s instructions; processed samples were read at a Luminex xMAP (BioRad) and analyzed with the software Bio-Plex Manager 6.0 (BioRad). Dynamic stimulation of iPSC-derived β cells was instead performed on an automated perifusion system (BioRep^®^ Perifusion V2.0.0). For each line, one hundred clusters were picked and stimulated with HEPES-buffered solution (125 mM NaCl, 5.9 mM KCl, 2.56 mM CaCl_2_, 1 mM MgCl_2_, 25 mM HEPES, 0.1% BSA, pH 7.4) supplemented as follows: 0.5 mM glucose, 11mM glucose plus 50µM IBMX (Gibco) or 30 mM KCl. Insulin content was quantified by ultrasensitive ELISA kit (Mercodia).

#### Bioinformatic analysis

Inference of the alternative splicing events potentially occurring after disruption of the ASS at c.316-2 (AG) of *WFS1* coding sequence was conducted following preliminary examination of the cryptic splice sites within the exon 4 sequence (ENSE00000701011) ± 150 bp flanking intronic regions. Analysis of the canonical splice sites, branch points, cis-elements (exonic splicing enhancers - ESE - and exonic splicing silencer - ESS) and trans-acting splicing factor sequences, was performed by the Human Splice Finder Pro (https://hsf.genomnis.com). To model the most thermodynamically favorite alternative splicing events following the c.316-1G>A transition the MaxEntScan framework (33) based on the maximum entropy principle was used. For the generation of alternative splicing isoform consensus sequences from deep sequencing raw data, the trimmed paired-end reads were aligned by STAR v2.5.3a to custom reference genome consisting of the wild type *WFS1* mRNA sequence as deposited in the RefSeq database (NM_006005.3) and the three alternative splicing isoform sequences as obtained by Sanger sequencing (c.316del, c.316-456del and c.316-460del). Due to higher probability of sequencing errors associated with read length, -outFilterMismatchNmax parameter was set to 10. Reads that did not align to any sequence of the custom reference genome were processed to identify other alternative splicing isoforms, if any, and used to generate the relative consensus sequence using EMBOSS’ Cons software (https://github.com/kimrutherford/EMBOSS). Mapping of paired-end reads to the custom reference genome was visualized with Integrative Genomics Viewer (IGV) v2.5. To infer the tridimensional (3D)-structure of Wolframin N-terminal domain, the entire aminoacidic sequence of the protein was used with Ginzu protocol (34) to predict the functional domains of the protein. The sequence of N-terminal domain (residues 1-402) was then submitted to Robetta server’s RoseTTAFold (https://robetta.bakerlab.org/), following *ab initio* modeling pipeline. The structures of mutated isoforms were generated employing Rosetta’s comparative modelling (35) and using the de novo modelled WT N-terminal structure as reference. The molecular dynamics simulations and conformational change analysis were performed by Gromacs v.2021.7 (36).

#### Statistical analysis

GraphPad Prism software (9.0.1 version) was employed to perform statistical analyses. Assuming the normal distribution and according to type of data, Student’s unpaired or paired t-test (one or two-tailed) was used for comparison between two groups, while to evaluate the effects of two independent variables on a dependent variable, two-way ANOVA with Dunn-Šídák correction for multiple comparison test was applied. Data are graphed as mean±standard error of the mean (SEM) or as mean±standard deviation (SD). Error bar meaning, number and characteristics of replicates, and statistical analysis are reported in the figure legends.

## Results

### Analysis of patient’s *WFS1* mutations

The patient did not show any notable clinical manifestation before T1D diagnosis at the age of 5.2 years, with the exception of a transient mild speech delay. At age 8.7, WS1 was diagnosed by the Next Generation Sequencing (NGS)-based screening for monogenic diabetes-driving mutations. Residual c-peptide at baseline screening was 0.1 ng/mL, with HbA1c of 6.7% and an insulin requirement of 0.28 U/kg/day. MRI revealed slight bilateral atrophy of brainstem and optic nerves, as extensively described in our previous paper (37). Moreover, she presented a pubertal delay and the analysis of hormonal values suggested an early stage primary ovarian insufficiency (38). DNA-sequencing patterns were compared to the sequence of *WFS1* transcript variant 1 in GenBank (NCBI Ref. Seq: NM_006005.3; Gene ID: 7466), leading to detection of the heterozygous mutations c.316-1G>A and c.757A>T (**Fig. 1a-b**), and other homozygous benign variations as reported in **Table 1**. The c.316-1G>A transition disrupts the acceptor (3’) splice site of exon 4 with uncertain outcome (**Fig. 1a**) and was classified as likely pathogenic, according to the American College of Medical Genetics and Genomics (ACMG) guidelines (39). The c.757A>T transversion was classified as pathogenic and located on the exon 7, where it gives rise to a premature termination codon (PTC) at the amino acid residue 253 (p.Lys253Ter), as shown in **Fig. 1b**. Sanger sequencing on the PCR products from genomic DNA isolated from the family members confirmed that the patient inherited both c.316-1G>A and c.757A>T mutations from the father and the mother, respectively (**Fig. 1c**). Since the parents have no symptoms related to the syndrome, we assumed that both mutations do not exert any dominant negative effect and were transmitted as an autosomal recessive trait. By querying the ClinVar archives of NCBI, the Human Gene Mutation Database and the Leiden Open Variation Database (LOVD) we found that splicing site variations of *WFS1* gene account for 2.9% of the total mutations and, among them, only 1 out of 3 involves an ASS or its proximal polypyrimidine (PY) tract. In **Fig. 1d** we display the genomic position of the mutations carried by our patient and we highlight the previously reported ASS variants mapping to the intervening sequence (IVS)-1, IVS-4 and IVS-7 of the gene, as well (40, 41). To infer the effects of disruption of canonical AG ASS, we evaluated the cryptic splice sites along the *WFS1* exon 4 sequence (ENSE00000701011) ± 150 bp flanking intronic regions, by using the sequence analysis tool of the Human Splice Finder (**Fig. 2a-c**) and we identified a total of 4 ASSs with an HSF score ≥ 75 (**Fig. 2a** and **ESM Table 4**). Among these, the acceptor splice motif located at 6288996 (CACTACCTGC<u>AG</u>TT; HSF Score: 81.29) might cause a 21 bp deletion (c.316_336del) that preserves the open reading frame (ORF) generating a protein lacking a 7 aa fragment (VGKHYLQ) with apparent molecular weight of 99.5 kDa. None of these alternative splice sites were confirmed using inhomogeneous Markov models as shown by plotting the MaxEntScores ≥ 4 (**Fig 2a** and **ESM Table 4**). Interestingly, we observed that the exonic splicing auxiliary sequences ratio (ESR) resulted significantly higher (>15) along the cryptic acceptor splice sequence starting at 6289115 (CTTGGCGGAC<u>AG</u>AA; HSF Score 78.06) (**Figure 2b**). This positive ESR ratio was suggestive of an increased activity of exon splicing enhancer proteins as the analysis displayed the presence at the 5’ of exon 4 of the recognition motifs of the SC35, SRp40 and 9G8 splicing factors (**Fig. 2b**). We also find a total of 6 branch points (BP score > 65) along the sequence comprising between 6288948 and 6289146 that could be used from the cryptic ASSs (**Fig 2c** and **ESM Table 4**). To infer which ASS could be selected by the spliceosome when the disruption of the wild type (WT) site occurs, we applied a method based on Maximum Entropy Principle (33). We expected the abolishment of the WT ASS would drive the splicing machinery to utilize the cryptic sequence at position 6288996. However, since the c.316-1G>A mutation determines the 1 bp-shifting of the canonical AG motif, according to the Maximum Entropy model, this ASS was thermodynamically favored when compared to the cryptic motif at 6288996 (**Fig. 2d**). Nevertheless, the generation of the 1 bp-shifted splice site at position c.317 dramatically alters the open reading frame (ORF) and results in a PTC. Anyhow, the maximum entropy-based probabilistic model suggested the exon 4 skipping as the most probable outcome (MaxEntScore > 6) causing ORF breakdown and PTC formation as well (**Fig. 2d**).

**Figure 1.**
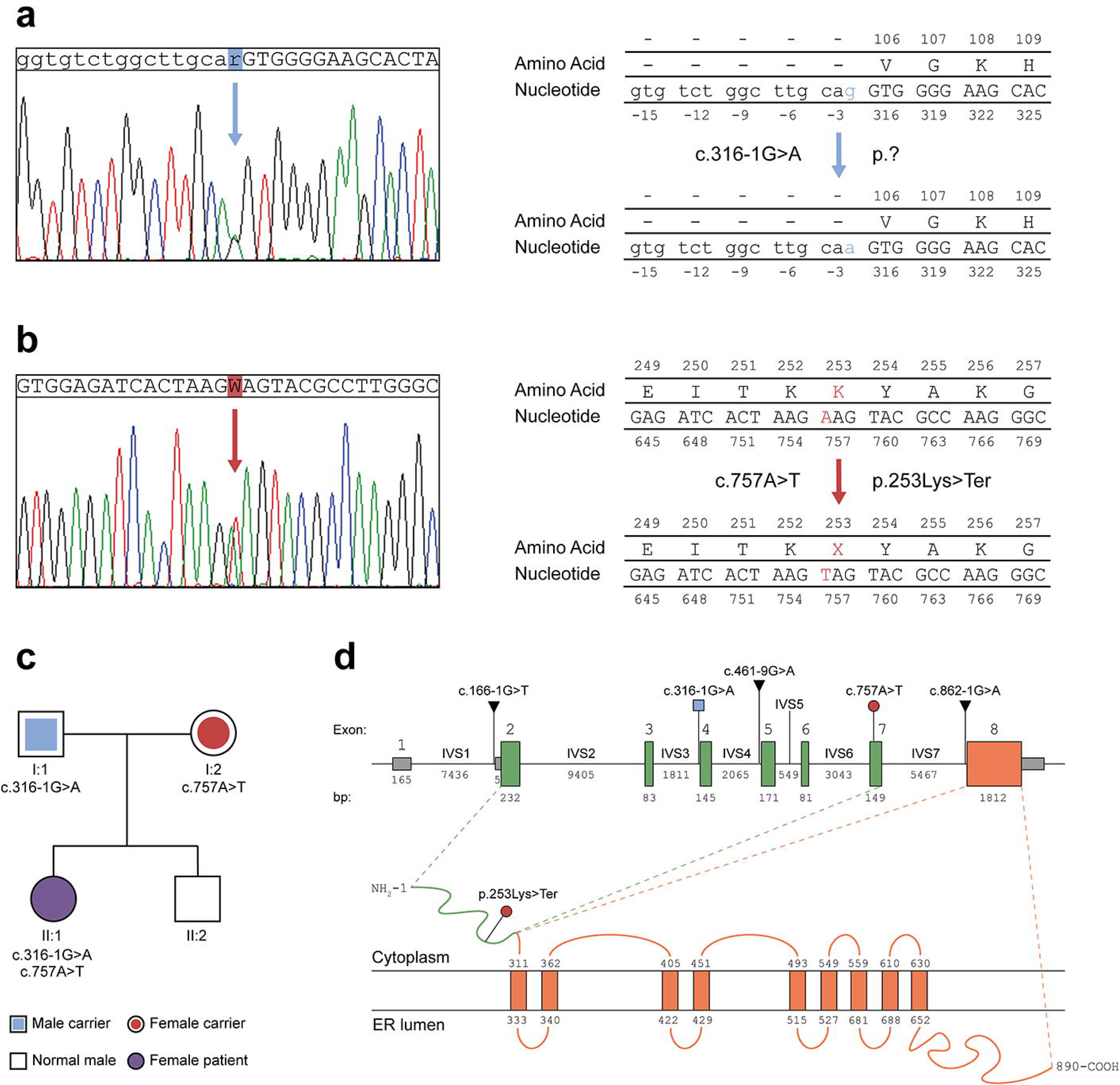
(**a-b**) Sanger sequencing chromatograms showing the heterozygous c.316-1G>A transition and the c.757A>T transversion in *WFS1* gene of the patient. The modified nucleotides are highlighted in blue and red, respectively, as well as the resulting aminoacid substitution in the protein sequence, if known. **(c)** Genetic pedigree of the patient’s family. The first line below each symbol represents generation and identification number. The heterozygous mutations found in the patient’s father and mother are reported; the patient’s brother carries no mutated allele. **(d)** Schematic representation of *WFS1* gene and Wolframin protein. The exons 1 (non-coding, grey), 2-7 (encoding for the N-terminal domain of wolframin, green boxes) and 8 (encoding for the transmembrane and C-terminal domains of wolframin, orange box) are reported. Genomic position of patient’s mutations are highlighted and color-coded as in panels A-B. The previously reported ASS variants are reported as well. The N-terminal domain (green) and transmembrane/C-terminal domains (orange) are shown along with the c.757A>T transversion introducing a premature stop codon.

**Figure 2.**
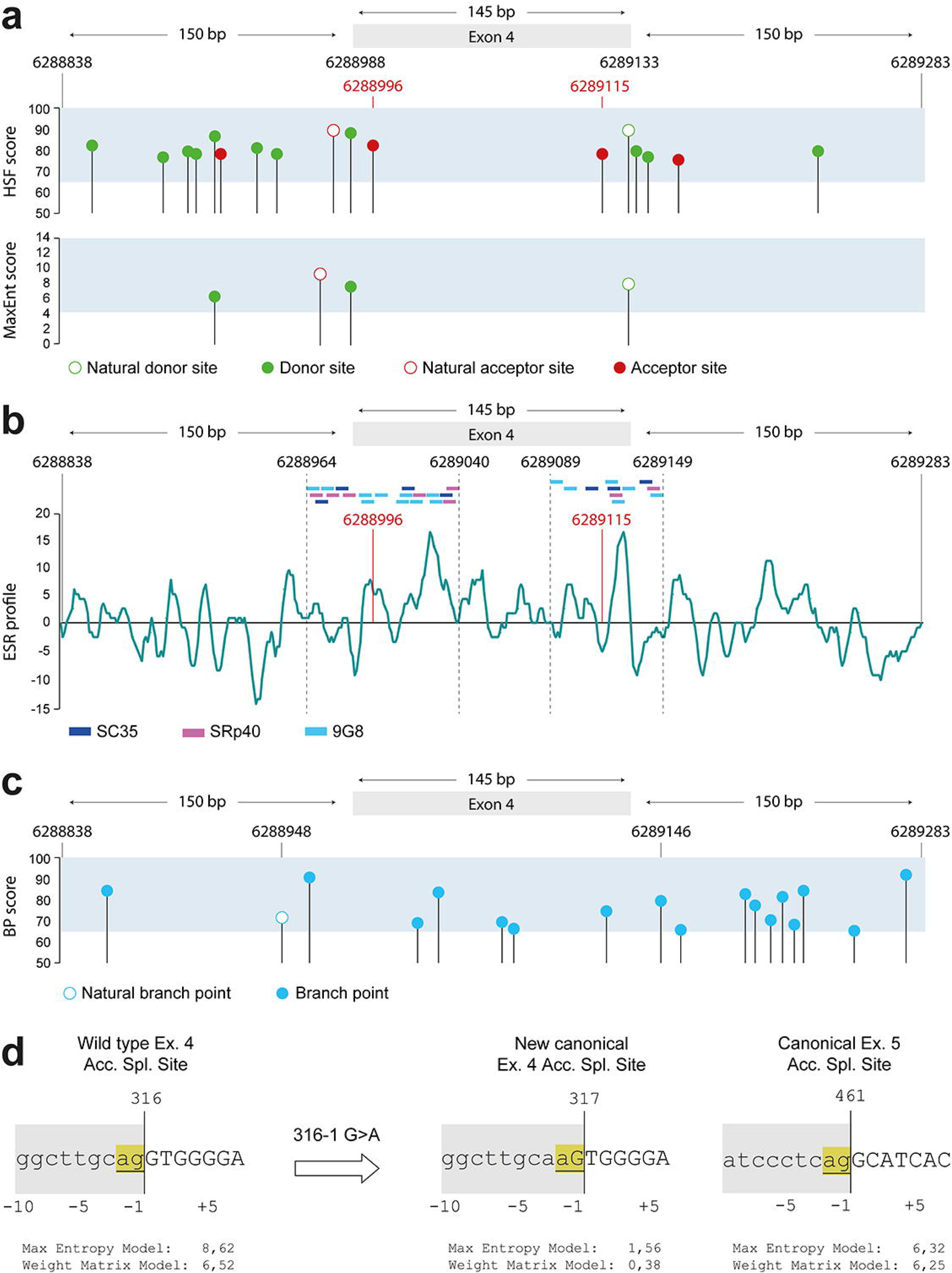
(**a**) Genomic positions of the natural and cryptic donor (green) and acceptor (red) splice sites along the *WFS1* exon 4 (145bp) and its flanking regions (150bp/each) as predicted by Human Splice Finder. Only the splice sites with an HSF score >75 and a MaxEnt score >4 are plotted. **(b)** Exonic splicing ratio (ESR) calculated as ratio between auxiliary splicing signals (ESE and ESS) along the same genomic region. The binding sites for the splicing factors SC35 (dark blue), SRp40 (purple) and 9G8 (light blue) are noted for the region of interest surrounding the cryptic ASSs inside exon 4. **(c)** Position of the natural and alternative branch points (light blue) along the analyzed genomic region. Only the branch points with a score >65 are plotted. The coordinates indicate the base number counting from the p-arm telomere of chromosome 4 (according to the UCSC Genome Browser on Human GRCh38/hg38 Assembly). **(d)** Results of maximum entropy modeling in presence of the wild type ASS and after the c.316-1G>A transition. The two most likely alternatives upon the canonical splice site disruption are reported. ASSs scored by the model are underlined and highlighted in yellow. Exonic bases are in upper case letters; intronic bases in lower case letters. The grey area represents the spliced-out sequence. Coordinates refer to the nucleotide position in the coding sequence.

### Characterization of the transcriptional outcomes from the c.316-1 G>A allele

To explore the transcriptional landscape arising from ASS disruption in IVS-3, we decided to exploit the stem cell technology. Specifically, we reprogrammed peripheral blood CD34^+^ cells of the patient into iPSCs, as previously described by our group (31), successfully generating a total of eight iPSC syngeneic clones with normal karyotype and conserved *WFS1* heterozygous mutations. Details on genomic integrity and analysis of pluripotency are reported in **ESM Fig. 1a-c**. We designed a PCR strategy to amplify *WFS1* transcripts along exons 3, 4 and 5, by using two pairs of primers as shown in **Fig. 3a**. When the *WFS1* mRNA between exon 3 and 5 was amplified in both WT and WFS1 iPSCs, we highlighted a potential mutation-related alternative splicing isoform as we observed an amplicon of ∼180bp in the WFS1 iPSCs (**Fig. 3b**). Both cell lines also display the expected amplicon of 324 bp, which derives from normal gene in the WT cells, and likely from the c.757A>T-carrying allele in the WFS1 ones. The amplification of the sequence between exons 3 and 4 or between exon 4 and 5 generated a single amplicon in WFS1 iPSCs, supporting the hypothesis that exon 4 skipping was the most probable event occurring in the c.316-1G>A-carrying pre-mRNA. (**Fig. 3b**). To determine the sequences of the observed PCR products, we TA-cloned the WFS1-derived 324bp- and 180bp-long bands and >20 clones were analyzed by Sanger sequencing. We found both natural and alternative splicing variants that we named according to Human Genome Variant Society nomenclature standard. Among the 324bp-long amplicons we identified two sequences: the WT (deriving from the c.757A>T-carrying allele) and the c.316del, corresponding to the maximum entropy-based prediction of alternative splicing resulting from the 1bp-shifting of ASS. Analyzing the 180bp-long amplicons we found two different sequences, one resulting from the skipping of the entire exon 4 (c.316_460del) and the other retaining the last four bases of the exon 4 (c.316_356del). Among the identified c.316-1G>A-carrying allele-deriving isoforms only the c.316_356del preserved the ORF (**Fig. 3c**). To exclude that reprogramming could alter the processing of *WFS1* pre-mRNA, as well as to rule out that observed isoforms were a specific characteristic of stem cells, due to a broader expression of splicing enhancer factors, we amplified the WT and WFS1 primary PBMC-derived cDNA by Ex3fw-Ex5rev primer pair, observing a complex pattern of amplicons in patient-derived PBMCs (**Fig. 3d**). Based on these findings, we generated a sequence library from both WT and WFS1-derived amplification products of the Ex3fw-Ex5rev primer pair and performed deep sequencing to identify all the potential alternative splicing isoforms occurring in the mutated cells. We performed paired-end sequencing on the PCR products from one WT subclone and two WFS1 clones. Assessment of read quality showed that R1 (forward) reads have poor quality after 210bp, while R2 (reverse) reads lose accuracy after 180bp. Preliminary STAR alignment to reference human genome hg38 revealed a reduced number of reads mapping to the *WFS1* gene in the affected cell-derived samples. This mirrored the percentage of trimmed paired-end reads per sample used to individuate the alternative splicing isoforms in WFS1 iPSCs (**Fig. 3e**). Interestingly, a total of 4 c.316-1G>A-carrying allele-deriving alternative splicing isoforms were identified in WFS1 iPSCs, whereas in WT cells the normal, correctly spliced transcript represented more than 99% of the mapped reads (**Fig. 3f**). NGS analysis thus confirmed sequences identified by cloning and Sanger sequencing and allowed to identify an additional splice variant: c.271_513del. Among detected transcripts, the c.316del and c.316_460del isoforms resulted in frameshift and PTC generation, whereas the c.316_356del and c.271_513del isoforms conserved the ORF, missing part of the *WFS1* mRNA. Specifically, c.316_356del lost the entire exon 4 with the exception of the last four nucleotides (AGAG), while c.271_513del included a larger deletion comprising the last 45 bp of exon 3, the entire exon 4 (145 bp) and the first 53 bp of exon 5 (**Fig. 3g** and **ESM Table 5**). Relative quantification of total amount of *WFS1* transcripts revealed a significant reduction in WFS1 iPSCs (p<0.05; one-tail t-test), but not the complete absence of stable transcripts (**Fig. 3h**). We speculated that stable low expression of *WFS1* total mRNAs in affected iPSCs might result from post-transcriptional regulation by non-sense mediated decay (NMD) of the transcript deriving from the c.757A>T-carrying allele and the 2 out of 4 transcripts deriving from the c.316-1G>A-carrying allele as well. Indeed, the two isoforms c.316del and c.316_460del were significantly upregulated by treatment with NMDI-14, a molecule that disrupts the SMG7-UPF1 interactions and strongly inhibits NMD (**Fig. 3i**).

**Figure 3.**
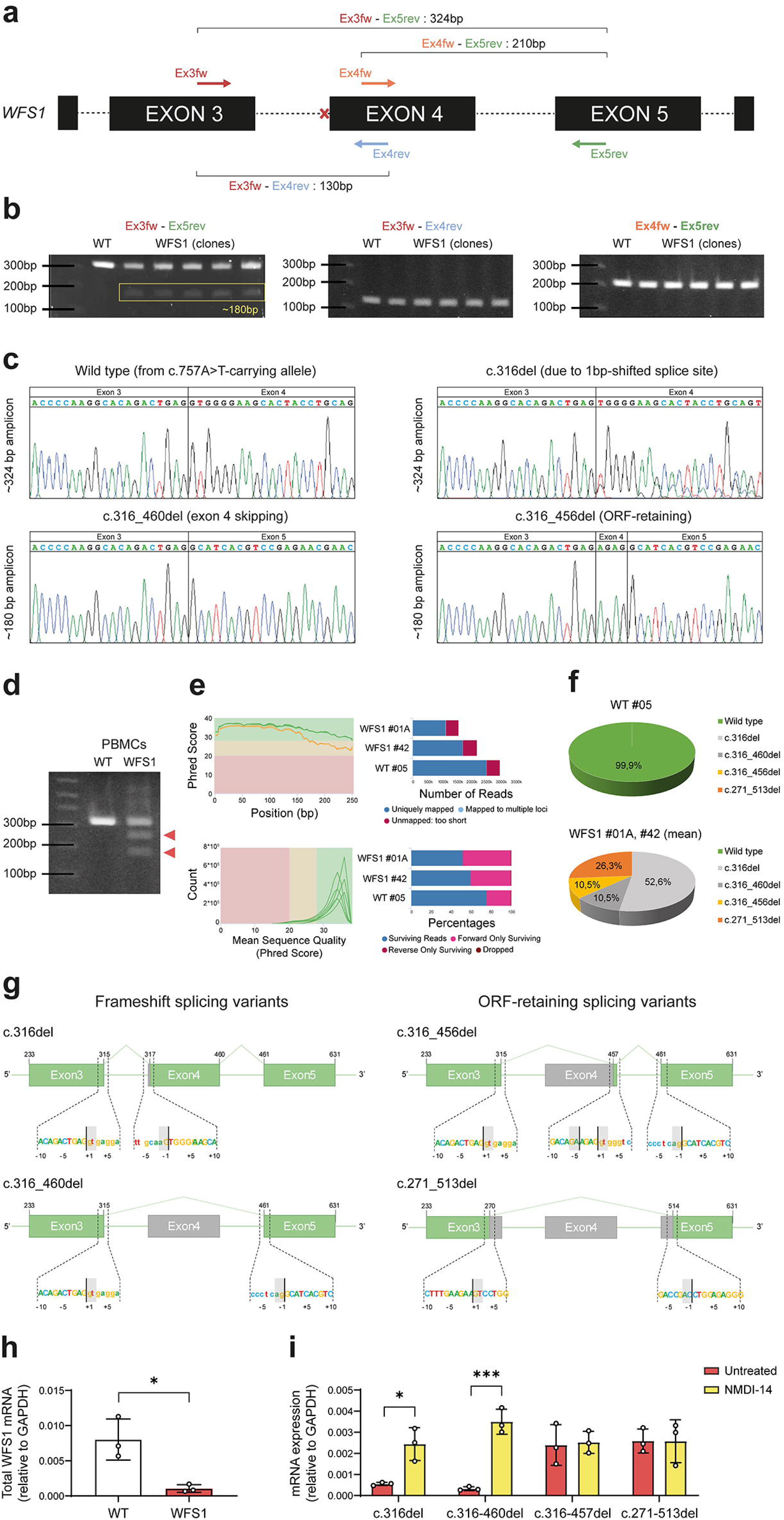
(**a**) Schematic representation of PCR strategy for the amplification of the WFS1 cDNA among exons 3, 4 and 5. Colored arrows show the mapping of the four primers along the WFS1 gene sequence. The red cross indicates the mutated canonical ASS. The theoretical length of PCR products obtained by performing amplification with different combinations of the indicated primers is reported. **(b)** Representative PCR results for the indicated primer pairs, performed on WT and five patient-derived clones. The red rectangle highlights the ∼180 bp-long amplicon obtained with the Ex3fw-Ex5rev primer pair. **(c)** Electropherograms from Sanger sequencing of WFS1-deriving PCR products. **(d)** Agarose gel electrophoresis of amplification with Ex3fw-Ex5rev from WT donor- and WFS1 patient-derived PBMCs. The additional amplicons, including the 180bp-length amplicon observed in WFS1 iPSCs, are indicated by red triangles. **(e)** High-throughput evaluation of RNA-seq data quality and goodness of alignment: *the sequence quality* graph, reporting the mean quality value (Phred score) across each base position in the R1 (green) and R2 (orange) reads; the *per sequence quality score* graph, reporting the number of reads with average quality scores. Red, yellow and green backgrounds highlight the Phred score ranges of bad, poor and good quality, respectively; *STAR alignment score*, reporting the number of mapped/unmapped reads per sample; percentage of reads per sample after adapter removal by Trimmomatic. **(f)** Percentage of NGS reads aligned to the wild type or the four alternative splice isoform consensus sequences in WT and WFS1 iPSCs. Alignment data for WFS1 are reported as mean of the two analyzed clones. **(g)** Schematic representation of the *WFS1* alternative splicing variants found by NGS, showing the conserved exonic sequence (green) and spliced-out sequence (grey). Sequences at the bottom of each structure zoom at the donor/acceptor alternative splice sites. **(h)** RT-qPCR of total *WFS1* mRNA, in WT and WFS1 iPSCs Data are plotted as mean±SD. N=3. The data were analyzed using the Student’s unpaired one-tailed t-test (*p<0.05). **(i)** RT-qPCR of the four *WFS1* isoforms, in WFS1 iPSCs after treatment with NMDI-14 or vehicle (untreated). Data are plotted as mean±SD. N=3 independent experiments. The data were analyzed using the two-way ANOVA with Dunn-Šídák correction (*p<0.05, ***p<0.001).

### Characterization of the translational outcomes from the c.316-1 G>A allele

The ORF-conserving c.316_356del and c.271_513del isoforms induce an in-frame skipping of 141bp and 243bp, resulting in 47aa (Val106-Arg152) and 81aa (Val91-Asp171) losses, respectively. This would lead to internally truncated forms of the protein, lacking part of the N-terminal domain but preserving transmembrane and C-terminal domains. To confirm the presence of mutated Wolframin, we performed immunoblot with two antibodies recognizing the N-terminal residues surrounding the Ala43 position and the Lys679-Phe783 C-terminal residues, respectively. The WT and WFS1-KO fibroblasts were used as controls of antibody specificity. As shown in **Fig. 4a**, the C-terminal-recognizing Ab confirmed that the Wolframin amount in the WFS1 iPSCs was ∼50% lower than the WT counterpart. This data highlighted that residual protein in affected cells derived from the c.316-1G>A-carrying allele only. Of note, the higher expression variability and the stable levels of Wolframin in WT and WFS1 iPSC clones, respectively, mirrored that observed at transcriptional level. As expected, the N-terminal-recognizing Ab did not detect altered isoforms in WFS1 iPSCs (**Fig. 4b**). These results confirm that at least one of the splicing isoforms found in WFS1 iPSCs leads to residual protein production, albeit missing a portion of the N-terminal domain. To uncover the impact of the portions lost upon alternative splicing events, we refined the structure of the N-terminal domain (residues 1-402) by following the *ab initio* modeling pipeline of RoseTTAFold. We deposited the *de novo* model in the ModelArchive Database (doi: 10.5452/ma-cg3qd) and used it to perform homology-based, comparative modeling of the mutated isoforms p.Val106-Arg152del and p.Val91-Thr170del (**Fig. 4c-d**). The p.Val106-Arg152del isoform partially preserved the WFS1-Calmodulin (CaM) binding site, whereas the p.Val91-Thr171 isoform lost the entire WFS1-CaM domain (42). Moreover, both isoforms resulted in altered N-terminal structure, potentially affecting interaction with other well-known WFS1 interactors, such as neuronal calcium sensor 1 (NCS1), that is involved in the mitochondria-associated membranes formation and calcium signaling (43). The dynamic simulation of predicted structures revealed higher instability of mutated N-terminal domain (> 0.5 Root Mean Standard Deviation, RMSD) compared to the WT ones (< 0.3 RMSD) after 10 ns into a salt-equilibrated aqueous solution (**Fig. 4e**). Finally, a radius of gyration of more than 30Å of the p.Val106-Arg152del isoform suggested a reduced compactness of its N-terminal domain (**Fig. 4f**). These *in silico* inferences support the hypothesis that the observed residual proteins in WFS1 cells might display reduced binding with cytoplasmic interactors as well as altered capacity to form homotetramers, which are considered the most stable and functional forms of Wolframin in the ER membrane (7, 8).

**Figure 4.**
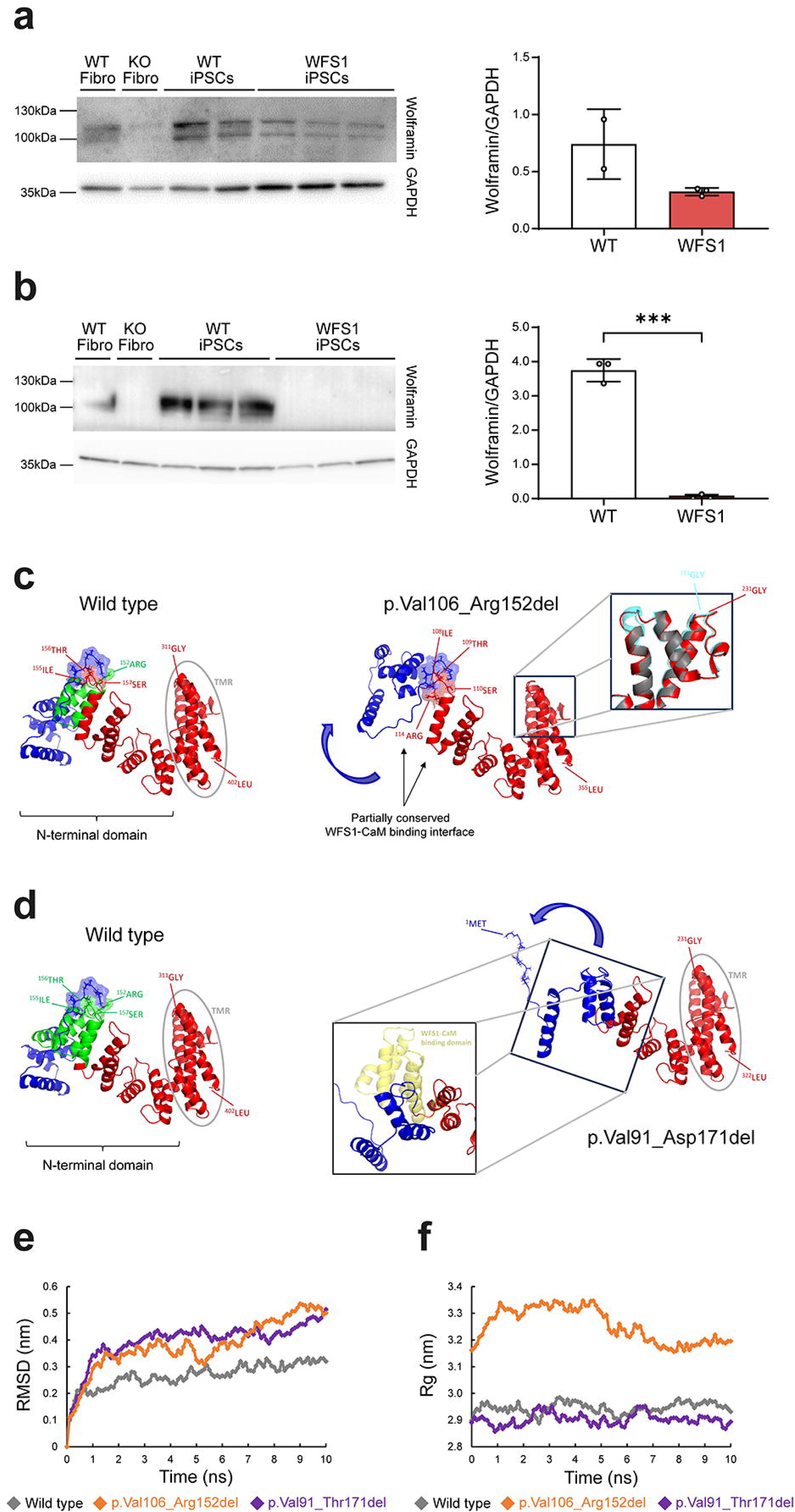
(**a-b**) Immunoblot analysis of Wolframin in WT and WFS1 iPSC clones recognized by polyclonal antibodies targeting the C-terminal (a) and N-terminal (b) domains. GAPDH was used as housekeeping gene. The densitometric analysis of the Wolframin/GAPDH ratio is reported. N=3. The data were analyzed using the Student’s unpaired two-tailed t-test (***p<0.001). **(c-d)** The three-dimensional models of WT (residues 1-402), and mutated p.Val106-Arg152del (residues 1-355) and p.Val91-Asp171del (residues 1-322) N-terminal domains, including the first two transmembrane regions (TMR). Proteins are represented as ribbons. Region lost in mutated isoforms (green), the mutation-induced conformational changes (blue) and the unaltered portion of the domain (red) are highlighted. In the pictures displaying the mutated isoforms, magnifications show the superimposed view of TMR alpha helices in WT (light blue) and p.Val106-Arg152del (red), and the superimposed view of WFS1-CaM binding domain (yellow) loss in p.Val91-Asp171del. In p.Val106-Arg152del, the partially conserved WFS1-CaM binding interface is shown as Connolly surface. **(e-f)** Results of molecular dynamics simulation, reporting RMSD of protein backbone atoms and radius of gyration (Rg) of the WT (grey) and mutated (orange/purple) N-terminal domains. Flattening of the RMSD and Rg plots of protein was observed around 10 ns at 300 K.

### CRISPR/Cas9-mediated gene correction of the c.316-1 G>A allele

To further confirm that the *WFS1* altered isoforms derive from the c.316-1G>A-carrying allele and to obtain a syngeneic allele-specific edited iPSC line to explore the functional impact of mutated transcripts, we performed a CRISPR/Cas9-mediated correction of the 316-1G>A point mutation. We designed single guide RNA (sgRNA) to target exon 4 of *WFS1* gene and trigger homologous-directed repair (HR) of the upstream intron-exon junction by using a single stranded oligonucleotide (ssODN) as shown in **Fig. 5a**. In the ssODN sequence we introduced A>G modification at c.316-1 to restore the natural ASS, and inserted four silent mutations at position p.113, 114, 115 and 116 (indicated in light blue in Figure 5a) to destroy the PAM sequence and the gRNA recognition motif, without altering aminoacidic code, to avoid re-targeting upon homologous recombination. Finally, we also introduced a SacI cut site (GAGCTC) to allow identification of gene engineered cells. Indeed, the gene edited clones were screened by amplifying the targeted sequence, followed by digestion with the SacI restriction enzyme (**Fig. 5b**). We obtained a total of six WFS1^wt/757A>T^ iPSC clones in which canonical splicing of *WFS1* mRNA and full-length isoform expression of Wolframin were restored (**Fig. 5c-d**); of note, protein levels were similar to those found in WT iPSCs (**Fig. 5c**). Restoring of the natural ASS was confirmed by RT-qPCR showing the absence of the 4 alternative splicing isoforms in WFS1^wt/757A>T^ iPSCs (**Fig. 5e**). All selected gene edited clones maintained normal morphology and karyotype, and displayed stemness properties as confirmed by expression of the pluripotency markers NANOG, OCT4 and SOX2 (**ESM Fig. 1d-f**).

**Figure 5.**
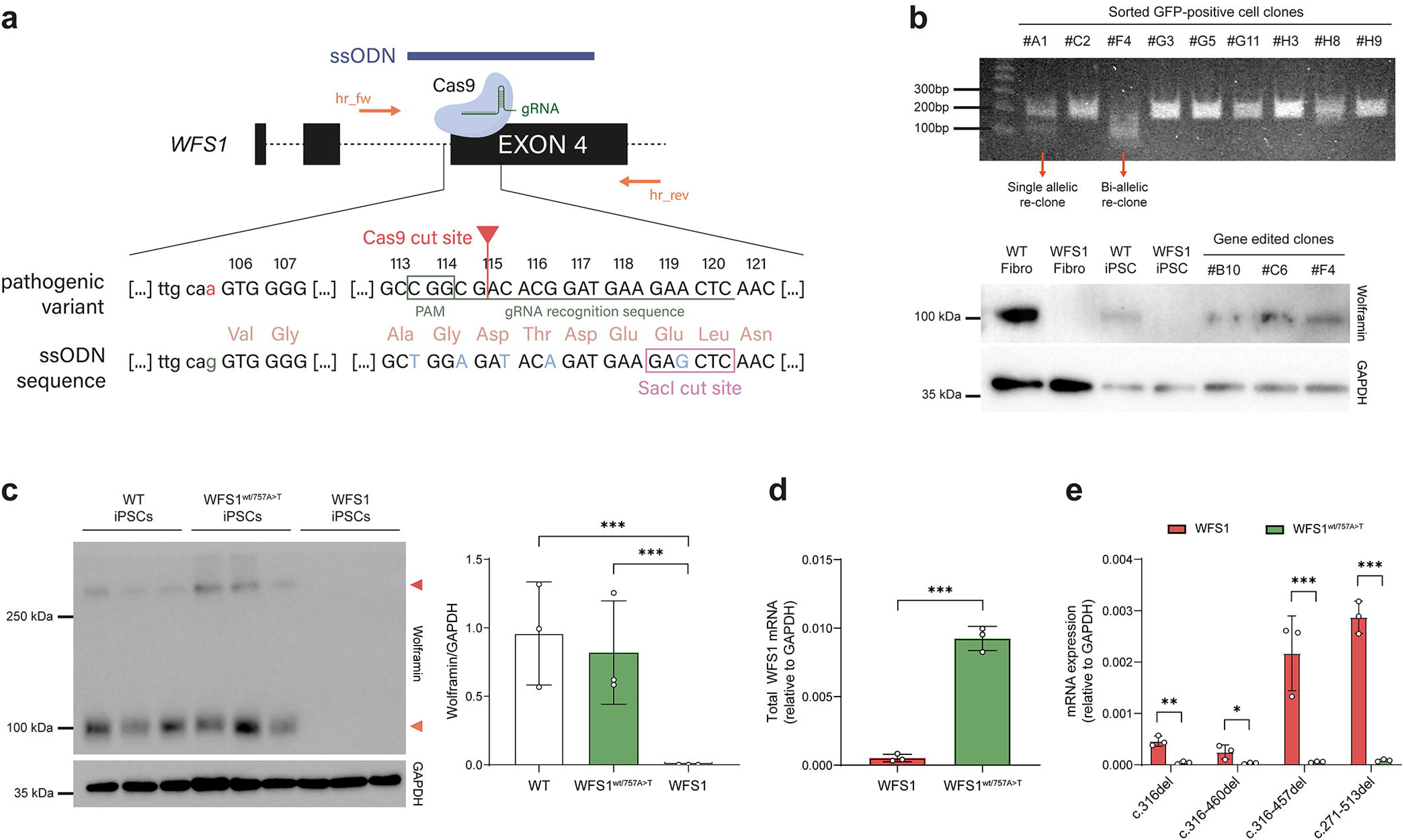
(**a**) Schematic representation of the c.316-1G>A point mutation correction strategy based on the Cas9/gRNA-mediated targeting of *WFS1* exon 4. Mapping of the ssODN (blue) and the fw/rev primer pair (dark/light orange) used for screening of the homology-directed repair is reported. Magnification highlights the comparison between the pathogenic variant containing the c.316-1G>A mutation with the ssODN sequence re-establishing the natural ASS upstream the exon 4. PAM sequence and gRNA recognition site are emphasized in dark green. Silent mutations and SacI cut site in ssODN sequence are reported in light blue and framed by pink rectangle, respectively. **(b)** Screening by SacI restriction enzyme digestion of GFP-positive iPSC clones and immunoblotting of Wolframin in WT and WFS1 Fibroblasts (as positive and negative controls, respectively), and WT, WFS1 and WFS1^wt/757A>T^ iPSC clones recognized by the polyclonal antibody targeting the N-terminal domain. GAPDH was used as housekeeping gene. (c) Immunoblot analysis of Wolframin expression in WT, WFS1wt/757A>T and WFS1 iPSCs. The relative quantification as Wolframin/GAPDH ratio is reported and expressed as mean±SD. N=3 different iPSC clones. The data were analyzed using the Student’s unpaired two-tailed t-test (***p<0.001). **(d)** RT-qPCR of total WFS1 mRNA and (e) of alternative splicing isoforms in WFS1 and WFS1^wt/757A>T^ iPSCs expressed as mean±SD. N=3. The data were analyzed using the two-way ANOVA with Dunn-Šídák correction (*p<0.05, **p<0.01, ***p<0.001).

### Expression of c.316-1 G>A allele-derived *WFS1* isoforms in insulin-producing β cells

Since pancreatic tissue and, in particular, insulin-producing β cells are affected in WS1, we aimed at exploring the impact of aberrant isoforms in the iPSC-derived β cells (iBeta). WFS1 and WFS1^wt/757A>T^ iPSCs were differentiated into pancreatic β cells following 24-days *in vitro* differentiation protocol (**ESM Fig. 2a**) and WFS1 cells did not display any alterations in the expression of key genes and proteins for the different developmental stages (**ESM Fig. 2b-c**). Furthermore, pancreatic differentiation efficiency and basal ghrelin, glucagon and insulin secretion did not change significantly among WT, WFS1 and WFS1^wt/757A>T^ cells at the iBeta stage (**ESM Fig. 2c-d**). Conversely, the WFS1 iBeta showed a slight, significant decrease of insulin secretion in response to high glucose (11 mM) plus the cAMP inducer IBMX at 15’, 20’ and 25’, in comparison to both WT and WFS1^wt/757A>T^ cells (**ESM Fig. 2e**). Given the pivotal role played by Wolframin in maintaining homeostasis and promoting insulin biosynthesis in pancreatic β cells (1), we expected WFS1 expression to increase during differentiation compared to the iPSC stage. Indeed, in WFS1^wt/757A>T^ cells we observed from 20 to 50-fold increase in *WFS1* mRNA levels after 24 days of differentiation, whereas the protein exceeded over 3-fold the GAPDH levels. Contrariwise, WFS1 cells did not display significant changes in both mRNA and protein levels after differentiation (**Fig. 6a-c**). We confirmed proper upregulation of *WFS1* upon pancreatic differentiation in the gene corrected clones by comparing the Wolframin levels with WT iBeta. Indeed, overall protein levels did not change between WT and WFS1^wt/757A>T^ iBeta (**ESM Fig. 3**). Moreover, by evaluating the levels of the c.316-1G>A-carrying allele-deriving mRNA isoforms we did not find any difference in c.316del and c.316_460del transcript expression between iPSCs and iBeta, confirming that such isoforms are subjected to NMD also in terminally differentiated cells. Surprisingly, we did not observe any increase in the ORF-maintaining isoforms after differentiation; conversely, WFS1 iBeta displayed a significant reduction of c.271_513del expression, suggesting that other mechanisms might be involved in their post-transcriptional regulation (**Fig. 6d**). As the Wolframin levels did not increase in the WFS1 iBeta, we hypothesized that the residual protein could not be able to guarantee the Wolframin-related functions in β cell stressogenic conditions. Finally, we observed the homotetrameric form of Wolframin in WFS1^wt/757A>T^ iBeta only, confirming that altered residual protein lacked the portion of the N-terminal domain involved in the tetramer formation in WFS1 cells (**Fig. 6a**).

**Figure 6.**
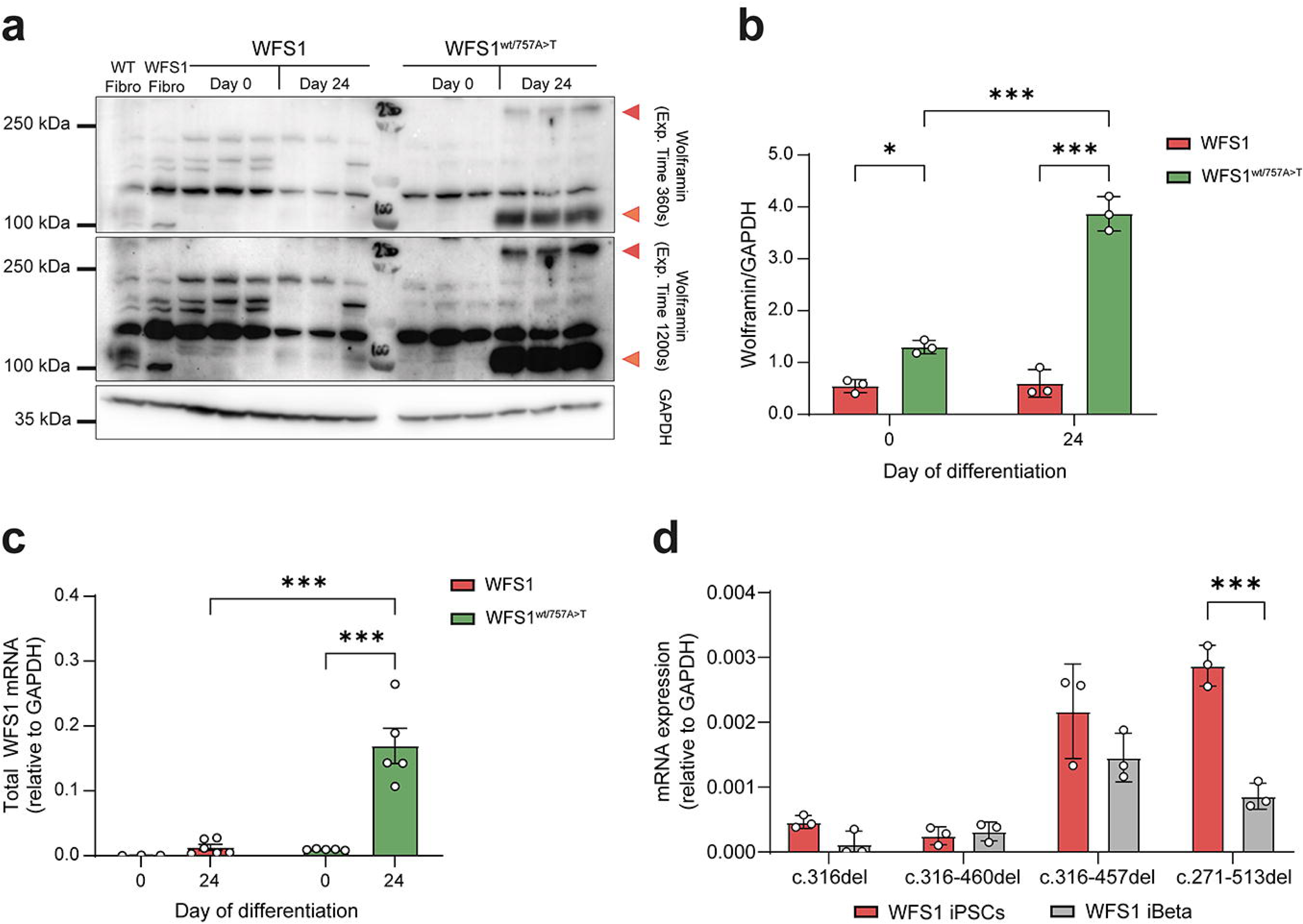
(**a**) Representative immunoblot showing Wolframin protein recognized by the anti-WFS1 C-terminal antibody before (day 0) and after (day 24) iPSCs differentiation into β cells. Monomeric form (100 kDa) and homotetrameric form (> 250 kDa) are indicated by orange and red triangles, respectively. Signal was acquired at different exposure times to detect homotetramers. **(b)** Relative quantification of Wolframin in WFS1 and WFS1^wt/757A>T^ iPSCs (day 0) and iBeta (day 24) expressed as mean±SD. N=3 independent in vitro differentiation experiments. The data were analyzed using the two-way ANOVA with Dunn-Šídák correction (* p<0.05, ***p<0.001). (c) RT-qPCR on total WFS1 mRNA in WFS1 and WFS1^wt/757A>T^ iPSCs (day 0) versus iBeta (day 24) reported as mean±SD. N=3-5 independent in vitro differentiation experiments. The data were analyzed using the two-way ANOVA with Dunn-Šídák correction (*** p<0.001). (d) Quantification of the alternative splicing isoforms in iPSCs and iBeta expressed as mean±SD. N=3. The data were analyzed using the two-way ANOVA with Dunn-Šídák correction (***p<0.001).

### Impact of *WFS1* aberrant isoforms on iBeta cell survival upon stress and inflammation

From a functional point of view, WS1 is characterized by aberrant ER stress responses and overall upregulation of ER stress-related genes in basal conditions (3). However, our data are in contrast with those reported in literature, as key ER stress-related markers were comparable in both WFS1 and WFS1^wt/757A>T^ iPSCs at proteins and transcriptional levels. Indeed, we did not find statistically significant differences in the protein levels of markers of the three ER stress-related pathways in WFS1 and WFS1^wt/757A>T^ iPSCs in basal conditions, with the exception of ATF4 and ERO1α (**ESM Fig. 4a**). More interestingly, no differences in the transcription levels of the ER stress-related genes *ATF4*, *BiP* and *CHOP* were detected after 2h, 4h, 8h, 16h or 24h upon TG treatment in WFS1 and WFS1^wt/757A>T^ iBeta, probably because residual protein activity is sufficient to control ER stress response (**ESM Fig. 4b**). Of note, WFS1 cells showed a significant decrease in sXBP-1/uXBP-1 ratio in comparison to the WFS1^wt/757A>T^ counterpart after 24h following TG treatment (**ESM Fig. 4b**), supporting the hypothesis that the IRE1α-dependent UPR may not be effective in controlling stress long term. However, reduced sXBP-1 levels did not correlate with higher apoptotic rate in WFS1 cells, supporting the hypothesis that cell death may be a consequence of ineffective short-term ER stress response. Although TG treatment induced the same cell death magnitude in WFS1 and WFS1^wt/757A>T^ iPSCs after 8h, we observed that recovery from TG-induced apoptosis occurred in the WFS1^wt/757A>T^ cells only. Indeed, the genetically corrected cells were able to reverse TG-induced apoptosis after 24h, while the percentage of Annexin-V-positive WFS1 cells remained around 40% up to 48h after stress induction (**ESM Fig. 4c**). We further examined *ATF4*, *BiP* and *CHOP* expression in WFS1 and WFS1^wt/757A>T^ iBeta 8h and 16h after TG treatment, confirming that TG stimulation induced a similar increase of UPR markers in both the iBeta cell lines at early time point. However, as observed in iPSCs, the UPR in WFS1 cells were less persistent over time compared to WFS1^wt/757A>T^ (**Fig. 7a, d, g**), as we reported a decrease of *ATF4* (0.24±0.11 at 8h vs 0.12±0.10 at 16h; p=0.12), *BiP* (1.46±0.48 at 8h vs 0.82±0.61 at 16h; p=0.41) and *CHOP* (0.35±0.11 at 8h vs 0.07±0.04 at 16h; p<0.01) in WFS1 iBeta after 16h. Both *ATF4* and *BiP* remained significantly higher compared to the baseline in WFS1^wt/757A>T^ cells at the same time point (over 5-fold for *ATF4* and over 2-fold for *BiP* after 16h; p<0.01). We then explored the impact of inflammation on the UPR in both WFS1 and WFS1^wt/757A>T^ iBeta. The exposure to inflammatory cytokines induced a significant UPR upregulation in WFS1^wt/757A>T^ iBeta, but not in WFS1 48h after treatment (**Fig. 7b-c, e-f, h-i**). Both TG and cytokines treatments triggered statistically significant higher rates of early and late apoptotic cell death in WFS1 compared to WFS1^wt/757A>T^ iBeta, as revealed by the frequency of Annexin V^+^ and P.I.^+^ cells, respectively (**Fig. 7j-m**). In particular, the WFS1 iBeta displayed a significant increase in Annexin-V^+^ cells after exposure to inflammatory cytokines and overall higher susceptibility to inflammatory signals compared to the gene edited counterpart. Of note, as it has been shown that both ER stress induction and inflammation are able to inhibit NMD and increase PTC-carrying mRNAs stability (44, 45), we wondered whether this might affect the PTC-carrying *WFS1* isoforms, resulting in toxic accumulation of the aberrant transcripts. Therefore, we evaluated the transcription levels of *UPF1* and *SMG7* genes, which encode for the main components of the NMD complex, and as expected, treatment with both TG and inflammatory cytokines significantly downregulated *SMG7* more than 2-fold compared to baseline, but not *UPF1*, in both WFS1 and WFS1^wt/757A>T^ iBeta (**Fig. 7n-o**). We further detected a dramatic increase of the PTC-carrying splicing isoforms c.316del and c.316_460del after exposure to cell stress inducer or pro-inflammatory cytokines (c.316del: >60-fold and 100-fold after 16h TG and 48h cytokines treatments, respectively; c.316_460del: >20-fold after either 16h TG or 48h cytokines treatments). These findings suggest that exogenous stress strongly affected the NMD-related gene *SMG7* and induced accumulation of PTC-carrying isoforms, correlating with increased cell death in WFS1 iBeta without UPR upregulation. A slight but not significant increase of the ORF-conserving isoforms c.316_356del and c.271_513del was detected as well (**Fig. 7p-q**).

**Figure 7.**
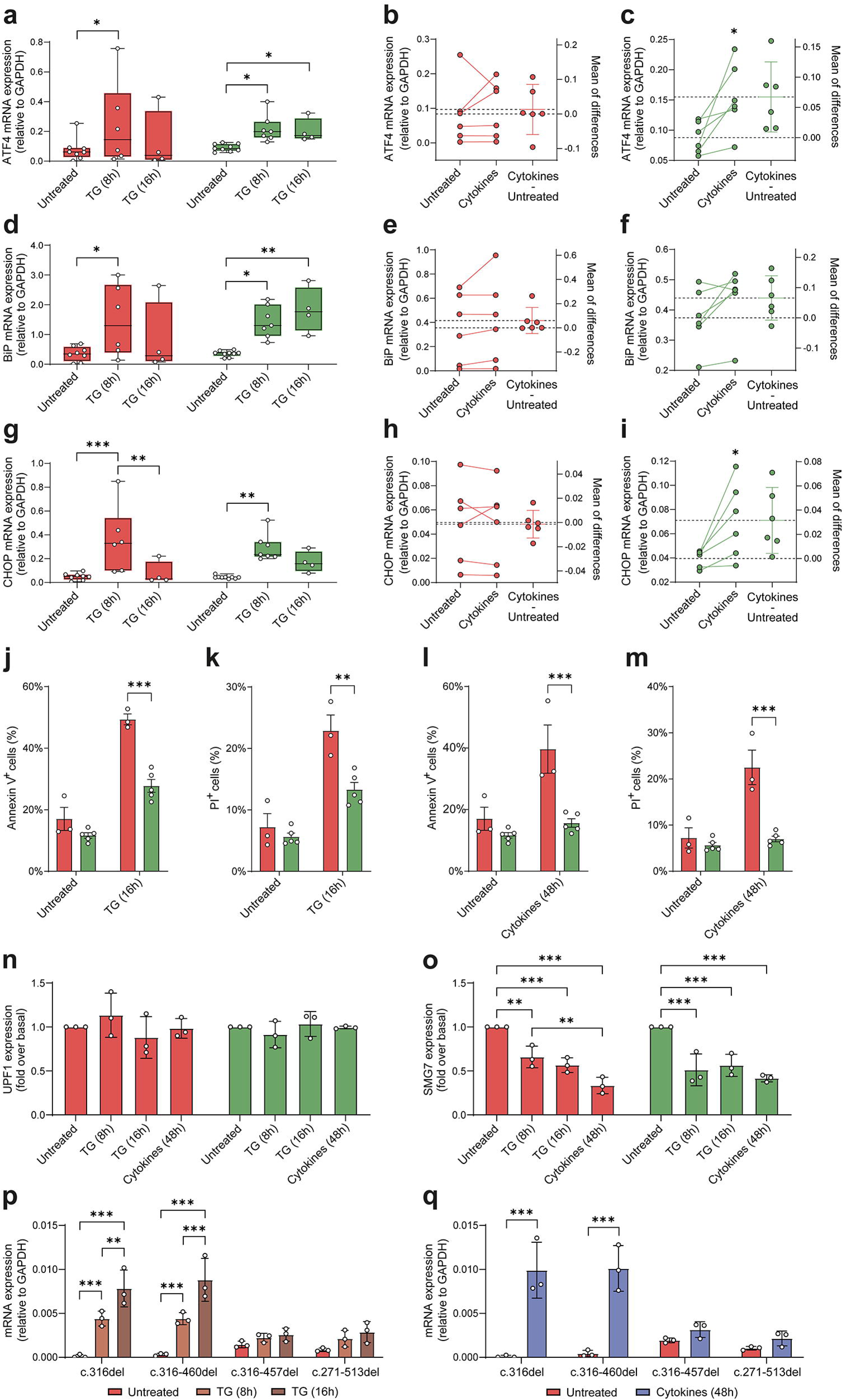
(**a**, **d, g)** Relative gene expression of ATF4, BiP and CHOP in WFS1 and WFS1^wt/757A>T^ β cells, after 8h or 16h post TG exposure. Data are plotted as mean±SEM. N=4-8 independent replicates. The data were analyzed using the two-way ANOVA with Dunn-Šídák correction (*p<0.05, **p<0.01). **(b-c, e-f, h-i)** Estimation plot including the mean of difference of ATF4, BiP and CHOP mRNAs calculated 48h after cytokines treatment (50 U/mL IL-1β, 1000 U/mL IFN-ɣ and 10 ng/mL TNF-α) in WFS1 (red) and WFS1^wt/757A>T^ (green) β cells. N=6 for WFS1, N=5 for WFS1^wt/757A>T^ iBeta. The data were analyzed using a paired Student’s t-test (*p<0.05). **(j-m)** Early and late apoptosis were measured as percentage (%) of Annexin V and P.I.-positive cells, respectively, in WFS1 and WFS1^wt/757A>T^ iBeta after 50 nM TG or inflammatory cytokines exposure. Data are plotted as mean±SD, N=3-5. The data were analyzed using the two-way ANOVA with Dunn-Šídák correction (**p<0.01, ***p<0.001) **(n-o)** Gene expression of the splicing factors UPF1 and SMG7 in WFS1 and WFS1^wt/757A>T^ iBeta upon TG or inflammatory cytokines treatment at the indicated times. Data are expressed as fold change over the untreated control. Data are expressed as mean±SD. N=3. Data were analyzed using the two-way ANOVA with Dunn-Šídák correction (**p<0.01, ***p<0.001). **(p)** Relative quantification of alternative splicing isoforms in WFS1 iBeta upon TG treatment for 8 and 16h. Data are plotted as mean±SD. N=3 independent experiments. Data were analyzed using the two-way ANOVA with Dunn-Šídák correction (**p<0.01, ***p<0.001). **(q)** Relative quantification of alternative splicing isoforms in WFS1 iBeta 48h upon inflammatory cytokines exposure. Data are plotted as mean±SD, N=3. The data were analyzed using the two-way ANOVA with Dunn-Šídák correction (***p<0.001).

## Discussion

Over the last few years, WS1 is assuming the connotation of a spectrum of disorders, due to the strong variability in terms of clinical manifestations and progression (46). Because of the role that Wolframin plays in a wide plethora of cellular processes, exploring the impact of *WFS1* variants on cell function and survival may represent a crucial step for their association with clinical phenotype.

Previous attempts to discover genotype and phenotype correlations led to the classification of *WFS1* mutations according to their impact on the protein expression and stability (16, 47, 48). A robust classification of *WFS1* genetic variants by disease onset and spectrum and severity of symptoms has been recently published as well (18). Studying the relationship between disease severity score reported by the Wolfram Unified Rating Scale and *WFS1* variants, the metanalysis performed by Urano’s group highlighted the importance of characterization of the classes of mutations for providing a more accurate prognoses of outcomes and a reliable therapy for patients with WS1.

The patient characterized in this work presents clinical symptoms with a mild progression of the disease in relation to age, with marginal optic nerve and hearing involvement (27), and residual circulating c-peptide (37). The mutation in exon 7 (c.757A>T) introduces a PTC; the other (c.316-1G>A) gives rise to 4 splice isoforms, of which 2 frameshift and 2 preserving the ORF. According to the classification offered by De Heredia (16), the introduction of a PTC before exon 8 causes NMD intervention, transcript degradation and therefore protein absence. By confirming that NMD induces complete degradation of PTC-carrying isoforms, we can state that both mutations c.757A>T and c.316-1G>A can be classified as Type 1 (16). However, the isoforms that preserve the ORF generate a functional residual protein, which would allow us to associate the patient genotype with an intermediate, mild-moderate phenotype, according to the classification reported by Urano’s work (18). In light of this consideration, the c.316-1G>A mutation has a transcriptional and translational phenotype more similar to that deriving from missense mutations, which maintain partial protein structure and, therefore, function.

The four novel isoforms of *WFS1* mRNA result from alternative splicing due to loss of the canonical ASS upstream the exon 4. In particular, 2 out of the 4 isoforms were correctly predicted by the use of a probabilistic model based on the maximum entropy principle. Indeed, we inferred that the two most probable alternative splicing isoforms could be c.316del (deriving from 1-bp shifting of the ASS following the mutation in c.316-1) and c.316_460del (losing of the exon 4 due to the direct skipping to the ASS of exon 5). Otherwise, the other 2 isoforms c.316_356del and c.271_513del were not predicted by the *in silico* analysis, but we found that the sequence surrounding the cryptic uncanonical splicing site generating the c.316_356del isoform resulted particularly enriched in ESEs. We excluded that the alternative splicing events occurring in the patient were a consequence of reprogramming of the somatic cells into iPSCs, refuting previous evidences reporting that PBMCs from the same patient displayed *WFS1* transcript originating from the c.757A>T-carrying allele only (28). Despite the design of primers to amplify *WFS1* mRNA was similar to that described by Panfili and colleagues, our protocol allows us to identify the alternative splicing isoforms deriving from the c.316-1G>A-carrying allele also in patient-derived primary PBMCs. Deep sequencing further confirmed the four isoforms, which we have chosen to classify based on the maintenance of the ORF and, therefore, on their ability to originate a protein, although with altered or reduced function.

Interestingly, the two ORF-maintaining isoforms c.316_356del and c.271_513del conserved the ability to produce a protein with intact transmembrane and C-terminal domains, thus potentially preserving its subcellular localization and intraluminal interactions. We confirmed expression in WFS1 iPSCs of Wolframin lacking part of N-terminal domain, but still maintaining the integrity of the C-terminal domain; this could explain the absence of significant alterations in ER stress markers upon ER stress induction and of the ATF6 cleaved form. However, we found that after differentiation into pancreatic β cells, the *WFS1* gene in affected cells was unable to reach the expression levels of the gene corrected clones. This resulted in low levels of residual proteins.

Our work pointed out another important aspect concerning the physiological mechanisms involved in pancreatic β cells: NMD. Indeed, the integrity and accuracy of transcript processing is pivotal for β cells to meet the physiological demands and pathophysiological challenges they face (21). The disruption of canonical splice sites and AS events introducing PTC may shift the balance towards the accumulation of aberrant transcripts during stress conditions, potentially affecting β cell survival (49).

We reported that the PTC-carrying c.316del and c.316_460del isoforms are regulated by NMD, as by inhibiting the UPF1/SMG7 interaction the degradation of such transcripts was prevented. Indeed, NMD orchestrates the Regulated Unproductive Splicing and Translation (RUST) mechanism to eliminate the PTC-containing transcript isoforms generated due to perturbed AS (50). Although the role of RUST in pancreatic β cells is still largely unknown, deregulation of NMD induced by islet stress or inflammation influences RUST, leading to accumulation of unproductive transcript isoforms, which are implicated in β cell dysfunction, vulnerability, and death (49). Interestingly, we found that patient-derived cells were particularly susceptible to the TG-induced cell death due to impairments to recover from Sarco-Endoplasmic Reticulum Calcium ATPase (SERCA) inhibition. In this context, we observed the early upregulation of UPR genes by both WFS1 and WFS1^wt/757A>T^ iBeta, but at later times (16h) WFS1 cells do not maintain the upregulation and, as a consequence, display increased apoptosis. A similar process seems to be involved in inflammatory cytokines-dependent apoptosis, which is higher in WFS1 iBeta compared to their genetically corrected counterpart. This result is driven by a lack of UPR upregulation upon cytokine exposure, which is, in contrast, efficiently achieved by WFS1^wt/757A>T^ iBeta. Our results point to lack of UPR upregulation as the cause of cell death following ER stress induction by inflammation, as already reported by other groups (51, 52). A previous study on the WS1 patient presented in this work reported a state of chronic inflammation associated with high levels of inflammatory cytokines, including IL-1β, TNF-α, IFN-γ and IL-6 (28). Increased secretion of inflammatory cytokines in the serum of the patient and susceptibility of *WFS1*-deficient β cells to exposure to pro-inflammatory factors could represent a new etiopathogenetic model for WS1-related diabetes. Both inflammation and other exogenous stress could be the cause of NMD inhibition and consequently of accumulation of aberrant transcripts, as shown by the increase of all alternative splicing isoforms upon stress induction. Indeed, after exogenous stimulus, we observed a decrease in *SMG7* levels and a critical increase of *WFS1* PTC-containing isoforms. Accumulation of these transcripts could be the cause of β cell dysfunction and apoptosis.

The case study we reported represents a new model to study Wolframin from a functional point of view, highlighting how each single mutation of *WFS1* gene can determine dramatically different outcomes from a clinical point of view. We underline the need for a better classification of mutations in the WFS1 gene, in order to explain the penetrance and expressivity of the clinical characteristics of the disease. Further understanding of mechanisms by which *WFS1* variants functionally affect ER stress or inflammatory responses may aid in the identification of other potential therapeutic targets.

Finally, the role of NMD in WS1 has not been considered until now, despite the majority of mutations in the *WFS1* gene introducing a PTC. Accumulation of *WFS1* unproductive transcripts whose translation into unfolded polypeptides overwhelms ER capacity may drive unresolved ER stress. The role of NMD among the different cell contexts may explain susceptibility to *WFS1* mutations of specific cell types. Although further research is needed to validate this hypothesis, NMD inhibition and PTC-carrying mRNA accumulation in WS1 could represent targetable mechanisms for the development of new therapeutic strategies to lengthen the half-life of the residual protein or compensate for missing functions. A similar rationale is highlighted in the cystic fibrosis field, another autosomal recessive disorder, where knowledge acquired from genotype-phenotype association studies has helped in the identification of effective therapeutic strategies and improved the prognosis (53–55).

### Ethics declarations

## Supporting information

Electronic supplementary material

## Acknowledgments

We thank the Advanced Light and Electron Microscopy BioImaging Center (ALEMBIC), at San Raffaele Scientific Institute, Milan (Italy), for confocal immunofluorescence images, and the Flow cytometry Resource, Advanced Cytometry Technical Applications Laboratory (FRACTAL), at San Raffaele Scientific Institute, Milan (Italy), for cell sorting experiments. We also thank Dr. Francesca Giannese and Dr. Dejan Lazarevic at Center for Omics Sciences (COSR) of San Raffaele Scientific Institute, Milan (Italy), for having provided support in library preparation, deep sequencing and bioinformatic analysis. We are grateful to Dr. Angelo Lombardo at the San Raffaele Telethon Institute for Gene Therapy (SR-TIGET), San Raffaele Scientific Institute, Milan (Italy), for the access to BLS2 work areas and the use of the 4D-Nucleofector System. Dr. Silvia Torchio conducted this study as partial fulfillment of an international PhD in Molecular Medicine at Vita-Salute San Raffaele University.

## Data availability

The *in silico* predicted N-terminal domain structure file of wild type WFS1 was deposited in the ModelArchive, together with procedures, ramachandran plots, inter-residue distance deviation and IDDT scores, and Gromacs configuration files (doi/10.5452/ma-cg3qd). The deep sequencing data as fastq files used to generate consensus sequences of alternative splicing isoforms of WFS1 are available in SRA database (BioProject PRJNA1109747). All raw data that were not directly included in the manuscript or that have not been deposited in online repositories, are available on request from the corresponding authors.

## Funding

This study was supported by a private family donation financing investigation on Wolfram Syndrome 1 at the Diabetes Research Institute (DRI) of the IRCCS San Raffaele Hospital. Part of the activities were also supported through the funds from the European Union - Next Generation EU - PNRR M6C2 - Investment 2.1 Enhancement and strengthening of NHS biomedical research (PNRR-MR1-2022- 12375914).

### Authors’ relationships and activities

The authors declare to have no conflicts of interest and that there are no relationships or activities that might bias, or be perceived to bias, their work.

### Contribution statement

RC, GF and LP were responsible for the conception and design of the study. ST contributed to the experimental design and to the collection and analysis of the data, together with RC and GS. RC, ST, GS, VZ, LM, MTL, FM and FC conducted wet experiments. SP supported collection of data during revision activities. RC performed bioinformatic analysis. RC, ST, GS and LP interpreted the results. PC followed genetic testing in clinical practice. Medical evaluations and patient specimen collection was performed by GF and RB. GR and VB supported cell reprogramming activities. VZ, LM, MTL, SP and VS performed stem cell differentiation and beta cell function analysis. RC and ST wrote the original draft. Critical review and editing of the manuscript was performed by RC, ST, GS, PC, GC, GF and LP. All authors approved the final version to be published. LP takes responsibility for the integrity of the data and is the guarantor of this work.

## Notes

https://www.ncbi.nlm.nih.gov/bioproject/?term=PRJNA1109747

https://modelarchive.org/doi/10.5452/ma-cg3qd

### Competing Interest Statement

The authors have declared no competing interest.

### Summary of Updates

The manuscript underwent significant changes following the revision process prior to publication. These changes included the addition of further evaluations and experimental sets, extensive rewriting to enhance readability and correct typographical errors, and adjustments to the figures, such as standardizing fonts and labels to improve legibility.

