## Supplementary material for "A *WFS1* variant disrupting acceptor splice site uncovers the impact of alternative splicing on β cell apoptosis in a patient with Wolfram syndrome": Electronic supplementary material

**ESM Table 1.** List of PCR primers and other oligonucleotides employed in the study.

| *Gene* (Protein) | Forward | Reverse |
|---|---|---|
| *ATF4* | GTTCTCCAGCGACAAGGCTA | ATCCTGCTTGCTGTTGTTGG |
| *HSPA5* (BiP) | TGTTCAACCAATTATCAGCAAACTC | TTCTGCTGTATCCTCTTCACCAGT |
| *DDIT3* (CHOP) | AGAACCAGGAAACGGAAACAGA | TCTCCTTCATGCGCTGCTTT |
| *GAPDH* | GTCTCCTCTGACTTCAACAGCG | ACCACCCTGTTGCTGTAGCCAA |
| *SMG7* | TTTCAGGAGGCAGTGGTGGATG | CAAACTCCTCTGGAAGTGGTGTC |
| *XBP-1* | CTGCCAGAGATCGAAAGAAGGC | CTCCTGGTTCTCAACTACAAGGC |
| *UPF1* | AACGAGCACCAAGGCATTGGCT | GGCTGCTTTGATAGTGCCTTCG |
| *c.316del* | CGACATTCTCCAGCAGCTC | CGACATTCTCCAGCAGCTC |
| *c.316_460del* | CCCAAGGCACAGACTGAGGC | CGACATTCTCCAGCAGCTC |
| *c.316_356del* | CTGGCCCTGGTGTTAGAGAC | TCGGACGTGATGCCTCTCTC |
| *c.271_513del* | AGCAGGAGAGGAGCGAAAG | CGCACGGCCCTCTCCAGGAC |
| *WFS1_ex3* | GGGCCTACAAAGGGAGACAT | - |
| *WFS1_ex4* | GGCGACACGGATTGAAGAACT | AGTTCTTCATCCGTGTCGCC |
| *WFS1_ex5* | - | CCAGTACATGACCAGGGCTG |
| *WFS1_gRNA* | TAATACGACTCACTATAGGAGTTCT TCATCCGTG | TTCTAGCTCTAAAACGCGACACGGAT GAAGAAC |
| *WFS1_hr* | CTTCCTCCTCACCCAGCCTG | GGGTCTTGGTCACTCACCTT |
| *WFS1_ssODN* | GAAAGGAGGTGGGCTGGCAGGGAGCATGGGGTGGGAGAGGGTCGGAGAAT CTGGAGGCTGACTGGTGTCTGGCTTGCAGGTGGGGAAGCACTACCTGCAGT TGGCTGGAGATACAGATGAAGAGCTCAACAGCTGCACCGCTGTGGACTGGC TGGTCCTCGCCGCGAAGCAGGGCCGTCGCGAGGCTGTGAAGCTGCTTC | |

**ESM Table 2.** List of TaqMan assays employed in the study.

| Gene ID | Company | Catalog Number |
|---|---|---|
| OCT4 (Hs00742896_s1) | Thermo Fisher Scientific | Cat#4331182 |
| NANOG (Hs04399610_g1) | Thermo Fisher Scientific | Cat#4331182 |
| SOX17 (Hs00751752_s1) | Thermo Fisher Scientific | Cat#4331182 |
| FOXA2 (Hs00232764_m1) | Thermo Fisher Scientific | Cat#4331182 |
| HNF1B (Hs00172123_m1) | Thermo Fisher Scientific | Cat#4331182 |
| PDX1 (Hs00236830_m1) | Thermo Fisher Scientific | Cat#4331182 |
| NGN3 (Hs01875204_s1) | Thermo Fisher Scientific | Cat#4331182 |
| NKX2.2 (Hs00159616_m1) | Thermo Fisher Scientific | Cat#4331182 |
| NKX6.1 (Hs00232355_m1) | Thermo Fisher Scientific | Cat#4331182 |
| GCG (Hs01031536_m1) | Thermo Fisher Scientific | Cat#4331182 |
| INS (Hs02741908_m1) | Thermo Fisher Scientific | Cat#4331182 |
| GAPDH (Hs99999905_m1) | Thermo Fisher Scientific | Cat#4331182 |

**ESM Table 3.** List of antibodies employed in the study and respective mode of use.

| Antibody | Species | Supplier | Code | Use | Dilution |
|---|---|---|---|---|---|
| Anti-GAPDH [6C5] | Mouse | Abcam | ab8245 | WB | 1:5000 |
| Alexa Fluor® 647 anti-Insulin [T56-706] | Mouse | BD | 565689 | FACS | 2µl x $10^6$ cells |
| Anti-Nanog | Rabbit | R&D | AF1997 | IF | 1:100 |
| PE anti-Nkx6.1 [R11-560] | Mouse | BD | 563023 | FACS | 5µl x $10^6$ cells |
| Anti-OCT4 [486] | Mouse | Novus | NBP2-15052 | IF | 1:100 |
| Alexa Fluor® 647 Anti-OCT3/4 [40] | Mouse | BD | 560329 | FACS | 4µl x $10^6$ cells |
| PE Anti-CXCR4 [12G5] | Mouse | BD | 557145 | FACS | 6µl x $10^6$ cells |
| Alexa Fluor® 488 anti-PDX1 [658A5] | Mouse | BD | 562274 | FACS | 5µl x $10^6$ cells |
| Anti-SSEA4 [MC813-70] | Mouse | Abcam | ab16287 | IF | 1:500 |
| Anti-Sox2 [245610] | Mouse | R&D | MAB2018 | IF | 1:100 |
| Anti-WFS1 C-terminal | Sheep | R&D | AF7417 | WB | 1:200 |
| Anti-WFS1 N-terminal | Rabbit | Cell Signalling | 8749 | WB | 1:1000 |
| Anti-ATF4 [EPR18111] | Rabbit | Abcam | ab184909 | WB | 1:1500 |
| Anti-ATF6 [1-7] | Mouse | Abcam | ab122897 | WB | 1:500 |
| Anti-BiP [C50B12] | Rabbit | Cell Signalling | #3177 | WB | 1:2000 |
| Anti-Calnexin [C5C9] | Rabbit | Cell Signalling | #2679 | WB | 1:1000 |

| | | | | | |
|---|---|---|---|---|---|
| Anti-phospho eIF2a [119A11] | Rabbit | Cell Signalling | #3597 | WB | 1:1000 |
| Anti-eIF2a | Mouse | Abcam | ab5369 | WB | 1:500 |
| Anti-ERO1α | Rabbit | Cell Signalling | #3264 | WB | 1:1000 |
| Anti-phospho IRE1a | Rabbit | Abcam | ab124945 | WB | 1:2000 |
| Anti-IRE1a [14C10] | Rabbit | Cell Signalling | #3294 | WB | 1:1000 |
| HRP-conjugated anti-mouse | Goat | R&D | HAF007 | WB | 1:1000 |
| HRP-conjugated anti-rabbit | Goat | R&D | HAF008 | WB | 1:1000 |
| HRP-conjugated anti-sheep | Donkey | R&D | HAF016 | WB | 1:1000 |
| Alexa Fluor™ Plus 488 anti-mouse | Goat | Thermo Fisher | A32723 | IF | 1:500 |
| Alexa Fluor™ Plus 488 anti-rabbit | Goat | Thermo Fisher | A32731 | IF | 1:500 |
| Alexa Fluor™ Plus 488 anti-goat | Donkey | Thermo Fisher | A32814 | IF | 1:500 |
| Alexa Fluor™ 546 anti-mouse | Goat | Thermo Fisher | A11030 | IF | 1:500 |
| Alexa Fluor™ 546 anti-goat | Rabbit | Thermo Fisher | A21085 | IF | 1:500 |

**ESM Table 4.** List of natural and cryptic donor and acceptor splicing sites, ESE/ESS signals and branch points along *WFS1* exon 4 and its 150bp-long flanking regions, as performed by using the Human Splice Finder Pro. Only HSF scores greater than 75 are reported.

| **HSF_Signals** | | | |
|---|---|---|---|
| **Type** | **Position** | **Motif** | **Value** |
| Donor splice site | 6288849 | CTGGTGTGA | 81.53 |
| Donor splice site | 6288851 | GGTGTGACC | 66.49 |
| Donor splice site | 6288886 | TGGGTGACA | 76.86 |
| Acceptor splice site | 6288886 | TGGGTGACAAAGGG | 69.99 |
| Donor splice site | 6288895 | AAGGGAAGT | 72.72 |
| Donor splice site | 6288899 | GAAGTGGGT | 78.76 |
| Acceptor splice site | 6288901 | AGTGGGTGAAAGGA | 68.74 |
| Donor splice site | 6288903 | TGGGTGAAA | 77.31 |
| Donor splice site | 6288913 | GAGGTGGGC | 85.92 |
| Acceptor splice site | 6288916 | GTGGGCTGGCAGGG | 77.91 |
| Acceptor splice site | 6288933 | ATGGGGTGGGAGAG | 66.2 |
| Donor splice site | 6288935 | GGGGTGGGA | 80.07 |
| Donor splice site | 6288945 | AGGGTCGGA | 77.67 |
| Acceptor splice site | 6288952 | GAGAATCTGGAGGC | 69.92 |
| Donor splice site | 6288969 | CTGGTGTCT | 74.19 |
| Donor splice site | 6288971 | GGTGTCTGG | 66.26 |
| (WT) Acceptor splice site | 6288975 | TCTGGCTTGCAGGT | 88.95 |
| Donor splice site | 6288984 | CAGGTGGGG | 87.37 |
| Acceptor splice site | 6288996 | CACTACCTGCAGTT | 81.29 |
| Donor splice site | 6289004 | GCAGTTGGC | 66.32 |
| Acceptor splice site | 6289018 | ACACGGATGAAGAA | 66.66 |
| Acceptor splice site | 6289028 | AGAACTCAACAGCT | 72.98 |
| Donor splice site | 6289059 | CTGGTCCTC | 67.66 |
| Acceptor splice site | 6289065 | CTCGCCGCGAAGCA | 71.7 |
| Acceptor splice site | 6289068 | GCCGCGAAGCAGGG | 72.8 |
| Acceptor splice site | 6289080 | GGCCGTCGCGAGGC | 68.89 |
| Acceptor splice site | 6289089 | GAGGCTGTGAAGCT | 68.12 |
| Donor splice site | 6289092 | GCTGTGAAG | 67.2 |
| Donor splice site | 6289109 | CCGGTGCTT | 73.55 |
| Acceptor splice site | 6289115 | CTTGGCGGACAGAA | 78.06 |
| (WT) Donor splice site | 6289129 | GAGGTGGGT | 88.85 |
| Donor splice site | 6289133 | TGGGTCTGT | 78.98 |
| Acceptor splice site | 6289135 | GGTCTGTGTGAGGC | 72.8 |
| Donor splice site | 6289137 | TCTGTGTGA | 69.1 |
| Donor splice site | 6289139 | TGTGTGAGG | 76.44 |
| Acceptor splice site | 6289141 | TGTGAGGCTTAGAA | 72.09 |
| Acceptor splice site | 6289146 | GGCTTAGAACAGCC | 73.12 |
| Acceptor splice site | 6289155 | CAGCCTCTGGAGGG | 75.12 |
| Acceptor splice site | 6289162 | TGGAGGGTTGAGCA | 66.07 |

| | | | |
|---|---|---|---|
| Donor splice site | 6289165 | AGGGTTGAG | 68.21 |
| Acceptor splice site | 6289165 | AGGGTTGAGCAGCT | 71.81 |
| Donor splice site | 6289177 | CTTGTAATG | 71.43 |
| Donor splice site | 6289228 | CTGGTGATG | 79.1 |
| Donor splice site | 6289236 | GCTGTTGGG | 66.22 |
| Acceptor splice site | 6289242 | GGGAAATTTCAGTT | 74.65 |
| Acceptor splice site | 6289256 | TCTGTTTTGCTGGT | 65.33 |
| Donor splice site | 6289265 | CTGGTGGCC | 71.26 |
| **MaxEnt_Signals** | | | |
| **Type** | **Position** | **Motif** | **Value** |
| Donor splice site | 6288913 | GAGGTGGGC | 6.32 |
| (WT) Acceptor splice site | 6288967 | GACTGGTGTCTGGCTTGCAGGTG | 8.62 |
| Donor splice site | 6288984 | CAGGTGGGG | 6.92 |
| (WT) Donor splice site | 6289129 | GAGGTGGGT | 7.07 |
| **BP_Signals** | | | |
| **Type** | **Position** | **Motif** | **Value** |
| Branch point | 6288857 | ACCCCAT | 82.59 |
| (WT) Branch point | 6288948 | GTCGGAG | 72.02 |
| Branch point | 6288963 | GGCTGAC | 88.62 |
| Branch point | 6289019 | CACGGAT | 68.98 |
| Branch point | 6289030 | AACTCAA | 82.03 |
| Branch point | 6289063 | TCCTCGC | 68.61 |
| Branch point | 6289070 | CGCGAAG | 65.54 |
| Branch point | 6289118 | GGCGGAC | 73.53 |
| Branch point | 6289146 | GGCTTAG | 79.00 |
| Branch point | 6289157 | GCCTCTG | 65.78 |
| Branch point | 6289191 | TGCTAAC | 81.83 |
| Branch point | 6289195 | AACTGAA | 77.09 |
| Branch point | 6289203 | AACTAAA | 70.20 |
| Branch point | 6289210 | ATCTTAC | 80.32 |
| Branch point | 6289215 | ACCAAAC | 68.60 |
| Branch point | 6289220 | ACCTAAC | 83.69 |
| Branch point | 6289247 | ATTTCAG | 65.58 |
| Branch point | 6289274 | TTCTCAT | 90.98 |
| **ESE_Signals** | | | |
| **Name** | **Position** | **Motif** | **CV** |
| ESE_9G8 (ESE Site) | 6288839 | GTGGAC | 77.79 |
| ESE_SRp40 (ESE Site) | 6288846 | TGCCTGG | 79.13 |
| ESE_9G8 (ESE Site) | 6288853 | TGTGAC | 60.94 |
| ESE_SC35 (ESE Site) | 6288856 | GACCCCAT | 77.21 |
| ESE_SC35 (ESE Site) | 6288861 | CATTTCTG | 75.30 |
| ESE_SRp40 (ESE Site) | 6288863 | TTTCTGC | 85.17 |
| ESE_SRp40 (ESE Site) | 6288877 | TTCCTGG | 82.18 |
| ESE_SC35 (ESE Site) | 6288882 | GGCCTGGG | 75.06 |
| ESE_9G8 (ESE Site) | 6288888 | GGTGAC | 82.69 |
| ESE_9G8 (ESE Site) | 6288891 | GACAAA | 66.58 |

| | | | |
|---|---|---|---|
| ESE_SRp40 (ESE Site) | 6288892 | ACAAAGG | 84.22 |
| ESE_ASF (ESE Site) | 6288893 | CAAAGGG | 79.12 |
| ESE_ASFB (ESE Site) | 6288893 | CAAAGGG | 79.15 |
| ESE_9G8 (ESE Site) | 6288899 | GAAGTG | 65.31 |
| ESE_9G8 (ESE Site) | 6288905 | GGTGAA | 70.61 |
| ESE_SRp40 (ESE Site) | 6288907 | TGAAAGG | 85.59 |
| ESE_ASF (ESE Site) | 6288908 | GAAAGGA | 74.17 |
| ESE_ASF (ESE Site) | 6288911 | AGGAGGT | 76.09 |
| ESE_9G8 (ESE Site) | 6288913 | GAGGTG | 60.47 |
| ESE_9G8 (ESE Site) | 6288916 | GTGGGC | 61.82 |
| ESE_SC35 (ESE Site) | 6288919 | GGCTGGCA | 80.41 |
| ESE_9G8 (ESE Site) | 6288920 | GCTGGC | 62.42 |
| ESE_SRp40 (ESE Site) | 6288922 | TGGCAGG | 79.37 |
| ESE_ASFB (ESE Site) | 6288923 | GGCAGGG | 72.61 |
| ESE_ASF (ESE Site) | 6288923 | GGCAGGG | 80.75 |
| ESE_9G8 (ESE Site) | 6288929 | GAGCAT | 60.14 |
| ESE_9G8 (ESE Site) | 6288941 | GGAGAG | 71.82 |
| ESE_ASF (ESE Site) | 6288942 | GAGAGGG | 81.92 |
| ESE_9G8 (ESE Site) | 6288944 | GAGGGT | 65.91 |
| ESE_SC35 (ESE Site) | 6288947 | GGTCGGAG | 78.75 |
| ESE_ASF (ESE Site) | 6288950 | CGGAGAA | 77.78 |
| ESE_ASFB (ESE Site) | 6288950 | CGGAGAA | 81.61 |
| ESE_9G8 (ESE Site) | 6288951 | GGAGAA | 74.84 |
| ESE_9G8 (ESE Site) | 6288952 | GAGAAT | 61.35 |
| ESE_9G8 (ESE Site) | 6288960 | GGAGGC | 70.94 |
| ESE_9G8 (ESE Site) | 6288961 | GAGGCT | 62.49 |
| ESE_9G8 (ESE Site) | 6288964 | GCTGAC | 78.39 |
| ESE_ASF (ESE Site) | 6288965 | CTGACTG | 75.80 |
| ESE_ASFB (ESE Site) | 6288965 | CTGACTG | 76.15 |
| ESE_SRp40 (ESE Site) | 6288966 | TGACTGG | 93.50 |
| ESE_SC35 (ESE Site) | 6288967 | GACTGGTG | 84.89 |
| ESE_9G8 (ESE Site) | 6288971 | GGTGTC | 63.29 |
| ESE_SRp40 (ESE Site) | 6288973 | TGTCTGG | 87.21 |
| ESE_SC35 (ESE Site) | 6288978 | GGCTTGCA | 81.20 |
| ESE_SRp40 (ESE Site) | 6288981 | TTGCAGG | 82.42 |
| ESE_ASFB (ESE Site) | 6288982 | TGCAGGT | 72.61 |
| ESE_ASF (ESE Site) | 6288982 | TGCAGGT | 74.99 |
| ESE_SRp55 (ESE Site) | 6288982 | TGCAGG | 75.89 |
| ESE_9G8 (ESE Site) | 6288989 | GGGGAA | 70.00 |
| ESE_9G8 (ESE Site) | 6288992 | GAAGCA | 68.33 |
| ESE_9G8 (ESE Site) | 6288996 | CACTAC | 59.33 |
| ESE_ASFB (ESE Site) | 6289002 | CTGCAGT | 71.77 |
| ESE_9G8 (ESE Site) | 6289007 | GTTGGC | 62.42 |
| ESE_SC35 (ESE Site) | 6289010 | GGCCGGCG | 84.59 |
| ESE_9G8 (ESE Site) | 6289011 | GCCGGC | 66.04 |
| ESE_ASFB (ESE Site) | 6289012 | CCGGCGA | 76.23 |

| | | | |
|---|---|---|---|
| ESE_9G8 (ESE Site) | 6289014 | GGCGAC | 86.31 |
| ESE_SRp40 (ESE Site) | 6289016 | CGACACG | 90.98 |
| ESE_ASFB (ESE Site) | 6289017 | GACACGG | 72.00 |
| ESE_ASF (ESE Site) | 6289017 | GACACGG | 86.93 |
| ESE_SRp55 (ESE Site) | 6289019 | CACGGA | 79.54 |
| ESE_ASF (ESE Site) | 6289021 | CGGATGA | 81.63 |
| ESE_ASFB (ESE Site) | 6289021 | CGGATGA | 83.00 |
| ESE_9G8 (ESE Site) | 6289023 | GATGAA | 83.49 |
| PESE (ESE Site) | 6289025 | TGAAGAAC | 62.05 |
| ESE_9G8 (ESE Site) | 6289026 | GAAGAA | 87.72 |
| ESE_Tra2 (ESE Site) | 6289027 | AAGAA | 100.34 |
| ESE_SC35 (ESE Site) | 6289029 | GAACTCAA | 77.46 |
| ESE_SRp40 (ESE Site) | 6289031 | ACTCAAC | 78.83 |
| ESE_SRp40 (ESE Site) | 6289034 | CAACAGC | 79.13 |
| ESE_Tra2 (ESE Site) | 6289035 | AACAG | 83.54 |
| ESE_SRp55 (ESE Site) | 6289041 | TGCACC | 75.76 |
| ESE_ASF (ESE Site) | 6289043 | CACCGCT | 74.06 |
| ESE_ASFB (ESE Site) | 6289043 | CACCGCT | 78.38 |
| ESE_SC35 (ESE Site) | 6289043 | CACCGCTG | 79.42 |
| ESE_ASF (ESE Site) | 6289048 | CTGTGGA | 74.58 |
| ESE_ASFB (ESE Site) | 6289048 | CTGTGGA | 77.00 |
| ESE_SRp55 (ESE Site) | 6289049 | TGTGGA | 78.83 |
| ESE_9G8 (ESE Site) | 6289050 | GTGGAC | 77.79 |
| ESE_SRp40 (ESE Site) | 6289052 | GGACTGG | 78.53 |
| ESE_SC35 (ESE Site) | 6289061 | GGTCCTCG | 81.57 |
| ESE_ASFB (ESE Site) | 6289067 | CGCCGCG | 79.08 |
| ESE_ASFB (ESE Site) | 6289069 | CCGCGAA | 71.00 |
| ESE_ASFB (ESE Site) | 6289072 | CGAAGCA | 72.00 |
| ESE_9G8 (ESE Site) | 6289073 | GAAGCA | 68.33 |
| ESE_ASF (ESE Site) | 6289075 | AGCAGGG | 75.34 |
| ESE_9G8 (ESE Site) | 6289077 | CAGGGC | 59.67 |
| ESE_ASF (ESE Site) | 6289079 | GGGCCGT | 79.30 |
| ESE_SC35 (ESE Site) | 6289080 | GGCCGTCG | 79.61 |
| ESE_9G8 (ESE Site) | 6289081 | GCCGTC | 62.62 |
| ESE_9G8 (ESE Site) | 6289089 | GAGGCT | 62.49 |
| ESE_9G8 (ESE Site) | 6289097 | GAAGCT | 67.32 |
| ESE_SRp55 (ESE Site) | 6289102 | TGCTTC | 81.58 |
| ESE_SC35 (ESE Site) | 6289107 | CGCCGGTG | 76.04 |
| ESE_ASF (ESE Site) | 6289107 | CGCCGGT | 86.40 |
| ESE_ASFB (ESE Site) | 6289107 | CGCCGGT | 93.31 |
| ESE_9G8 (ESE Site) | 6289119 | GCGGAC | 77.79 |
| ESE_ASF (ESE Site) | 6289120 | CGGACAG | 77.08 |
| ESE_ASFB (ESE Site) | 6289120 | CGGACAG | 79.92 |
| ESE_SC35 (ESE Site) | 6289121 | GGACAGAA | 75.86 |
| ESE_SRp40 (ESE Site) | 6289123 | ACAGAAG | 79.90 |
| ESE_ASFB (ESE Site) | 6289124 | CAGAAGA | 80.00 |

| | | | |
|---|---|---|---|
| ESE_ASF (ESE Site) | 6289124 | CAGAAGA | 84.25 |
| ESE_9G8 (ESE Site) | 6289126 | GAAGAG | 84.70 |
| ESE_ASF (ESE Site) | 6289127 | AAGAGGT | 78.72 |
| ESE_Tra2 (ESE Site) | 6289127 | AAGAG | 89.41 |
| ESE_9G8 (ESE Site) | 6289129 | GAGGTG | 60.47 |
| ESE_SRp55 (ESE Site) | 6289133 | TGGGTC | 76.72 |
| ESE_SC35 (ESE Site) | 6289135 | GGTCTGTG | 85.81 |
| ESE_SRp55 (ESE Site) | 6289139 | TGTGTG | 75.05 |
| ESE_SRp40 (ESE Site) | 6289141 | TGTGAGG | 78.41 |
| ESE_SRp55 (ESE Site) | 6289143 | TGAGGC | 77.42 |
| ESE_9G8 (ESE Site) | 6289144 | GAGGCT | 62.49 |
| ESE_SRp40 (ESE Site) | 6289149 | TTAGAAC | 79.25 |
| ESE_9G8 (ESE Site) | 6289152 | GAACAG | 62.96 |
| ESE_Tra2 (ESE Site) | 6289153 | AACAG | 83.54 |
| ESE_SC35 (ESE Site) | 6289156 | AGCCTCTG | 84.15 |
| ESE_SRp40 (ESE Site) | 6289158 | CCTCTGG | 89.96 |
| ESE_ASF (ESE Site) | 6289159 | CTCTGGA | 76.04 |
| ESE_ASFB (ESE Site) | 6289159 | CTCTGGA | 79.92 |
| ESE_9G8 (ESE Site) | 6289164 | GAGGGT | 65.91 |
| ESE_9G8 (ESE Site) | 6289168 | GTTGAG | 63.29 |
| ESE_SRp55 (ESE Site) | 6289172 | AGCAGC | 75.18 |
| ESE_SRp55 (ESE Site) | 6289184 | TGCTGC | 76.40 |
| ESE_9G8 (ESE Site) | 6289199 | GAACAA | 65.98 |
| ESE_Tra2 (ESE Site) | 6289200 | AACAA | 94.47 |
| ESE_Tra2 (ESE Site) | 6289207 | AAAAT | 81.36 |
| ESE_SRp40 (ESE Site) | 6289216 | CCAAACC | 81.40 |
| ESE_SRp40 (ESE Site) | 6289221 | CCTAACG | 80.20 |
| ESE_ASF (ESE Site) | 6289226 | CGCTGGT | 77.90 |
| ESE_ASFB (ESE Site) | 6289226 | CGCTGGT | 83.69 |
| ESE_9G8 (ESE Site) | 6289230 | GGTGAT | 69.60 |
| ESE_9G8 (ESE Site) | 6289233 | GATGCT | 63.09 |
| ESE_SC35 (ESE Site) | 6289252 | AGTTTCTG | 79.24 |
| ESE_SRp40 (ESE Site) | 6289262 | TTGCTGG | 80.92 |
| ESE_9G8 (ESE Site) | 6289267 | GGTGGC | 66.72 |
| **ESS_Signals** | | | |
| **Name** | **Position** | **Motif** | **CV** |
| ESS_hnRNPA1 (ESS Site) | 6288837 | TAGTGG | 82.27 |
| Sironi_motif2 (ESS Site) | 6288838 | AGTGGAC | 66.73 |
| Sironi_motif2 (ESS Site) | 6288850 | TGGTGTG | 63.08 |
| Sironi_motif2 (ESS Site) | 6288851 | GGTGTGA | 69.67 |
| Sironi_motif3 (ESS Site) | 6288855 | TGACCCCA | 64.15 |
| Sironi_motif3 (ESS Site) | 6288856 | GACCCCAT | 61.86 |
| Sironi_motif3 (ESS Site) | 6288865 | TCTGCCCC | 73.33 |
| Sironi_motif3 (ESS Site) | 6288871 | CCTTCCTT | 65.62 |
| Sironi_motif3 (ESS Site) | 6288875 | CCTTCCTG | 68.80 |
| Sironi_motif2 (ESS Site) | 6288883 | GCCTGGG | 60.01 |

| | | | |
|---|---|---|---|
| Sironi_motif2 (ESS Site) | 6288884 | CCTGGGT | 61.65 |
| Sironi_motif2 (ESS Site) | 6288886 | TGGGTGA | 60.21 |
| Sironi_motif1 (ESS Site) | 6288892 | ACAAAGGG | 64.83 |
| Sironi_motif2 (ESS Site) | 6288894 | AAAGGGA | 62.87 |
| ESS_hnRNPA1 (ESS Site) | 6288894 | AAAGGG | 65.13 |
| ESS_hnRNPA1 (ESS Site) | 6288895 | AAGGGA | 85.13 |
| Sironi_motif2 (ESS Site) | 6288898 | GGAAGTG | 62.31 |
| Sironi_motif2 (ESS Site) | 6288899 | GAAGTGG | 60.55 |
| Sironi_motif1 (ESS Site) | 6288899 | GAAGTGGG | 65.72 |
| ESS_hnRNPA1 (ESS Site) | 6288900 | AAGTGG | 67.03 |
| Sironi_motif2 (ESS Site) | 6288901 | AGTGGGT | 81.60 |
| Sironi_motif2 (ESS Site) | 6288903 | TGGGTGA | 60.21 |
| Sironi_motif1 (ESS Site) | 6288907 | TGAAAGGA | 62.38 |
| Sironi_motif2 (ESS Site) | 6288907 | TGAAAGG | 65.39 |
| ESS_hnRNPA1 (ESS Site) | 6288909 | AAAGGA | 67.98 |
| ESS_hnRNPA1 (ESS Site) | 6288910 | AAGGAG | 73.94 |
| Sironi_motif1 (ESS Site) | 6288910 | AAGGAGGT | 76.76 |
| Sironi_motif2 (ESS Site) | 6288912 | GGAGGTG | 77.46 |
| ESS_hnRNPA1 (ESS Site) | 6288913 | GAGGTG | 74.41 |
| Sironi_motif2 (ESS Site) | 6288914 | AGGTGGG | 73.16 |
| Sironi_motif2 (ESS Site) | 6288915 | GGTGGGC | 87.38 |
| Sironi_motif2 (ESS Site) | 6288919 | GGCTGGC | 66.88 |
| Sironi_motif2 (ESS Site) | 6288923 | GGCAGGG | 69.72 |
| Sironi_motif2 (ESS Site) | 6288924 | GCAGGGA | 66.62 |
| ESS_hnRNPA1 (ESS Site) | 6288925 | CAGGGA | 85.13 |
| Sironi_motif2 (ESS Site) | 6288931 | GCATGGG | 69.50 |
| Sironi_motif1 (ESS Site) | 6288932 | CATGGGGT | 67.99 |
| Sironi_motif2 (ESS Site) | 6288932 | CATGGGG | 73.71 |
| Sironi_motif2 (ESS Site) | 6288934 | TGGGGTG | 71.04 |
| Sironi_motif2 (ESS Site) | 6288935 | GGGGTGG | 67.96 |
| Sironi_motif2 (ESS Site) | 6288936 | GGGTGGG | 76.91 |
| Sironi_motif2 (ESS Site) | 6288937 | GGTGGGA | 86.58 |
| Sironi_motif2 (ESS Site) | 6288939 | TGGGAGA | 60.21 |
| Sironi_motif2 (ESS Site) | 6288941 | GGAGAGG | 77.46 |
| Sironi_motif1 (ESS Site) | 6288941 | GGAGAGGG | 79.52 |
| ESS_hnRNPA1 (ESS Site) | 6288942 | GAGAGG | 74.65 |
| Sironi_motif2 (ESS Site) | 6288943 | AGAGGGT | 78.55 |
| ESS_hnRNPA1 (ESS Site) | 6288944 | GAGGGT | 87.98 |
| Sironi_motif2 (ESS Site) | 6288946 | GGGTCGG | 60.01 |
| Sironi_motif2 (ESS Site) | 6288947 | GGTCGGA | 71.43 |
| ESS_hnRNPA1 (ESS Site) | 6288952 | GAGAAT | 68.22 |
| Sironi_motif2 (ESS Site) | 6288957 | TCTGGAG | 66.68 |
| Sironi_motif1 (ESS Site) | 6288958 | CTGGAGGC | 78.22 |
| Sironi_motif2 (ESS Site) | 6288959 | TGGAGGC | 62.77 |
| Sironi_motif2 (ESS Site) | 6288960 | GGAGGCT | 65.40 |
| ESS_hnRNPA1 (ESS Site) | 6288961 | GAGGCT | 77.98 |

| | | | |
|---|---|---|---|
| Sironi_motif2 (ESS Site) | 6288966 | TGACTGG | 65.39 |
| Sironi_motif2 (ESS Site) | 6288973 | TGTCTGG | 68.44 |
| Sironi_motif2 (ESS Site) | 6288982 | TGCAGGT | 60.74 |
| Sironi_motif2 (ESS Site) | 6288983 | GCAGGTG | 60.55 |
| ESS_hnRNPA1 (ESS Site) | 6288984 | CAGGTG | 70.60 |
| Sironi_motif2 (ESS Site) | 6288985 | AGGTGGG | 73.16 |
| Sironi_motif2 (ESS Site) | 6288986 | GGTGGGG | 97.41 |
| Sironi_motif2 (ESS Site) | 6288988 | TGGGGAA | 60.21 |
| Sironi_motif3 (ESS Site) | 6288997 | ACTACCTG | 65.95 |
| Sironi_motif2 (ESS Site) | 6289006 | AGTTGGC | 75.67 |
| ESS_hnRNPA1 (ESS Site) | 6289019 | CACGGA | 65.84 |
| Sironi_motif1 (ESS Site) | 6289026 | GAAGAACT | 63.88 |
| Sironi_motif1 (ESS Site) | 6289034 | CAACAGCT | 73.34 |
| Sironi_motif2 (ESS Site) | 6289047 | GCTGTGG | 63.60 |
| Sironi_motif2 (ESS Site) | 6289049 | TGTGGAC | 73.56 |
| Sironi_motif2 (ESS Site) | 6289052 | GGACTGG | 62.31 |
| Sironi_motif2 (ESS Site) | 6289057 | GGCTGGT | 64.85 |
| Sironi_motif3 (ESS Site) | 6289064 | CCTCGCCG | 67.10 |
| Sironi_motif2 (ESS Site) | 6289075 | AGCAGGG | 65.97 |
| Sironi_motif2 (ESS Site) | 6289076 | GCAGGGC | 67.43 |
| ESS_hnRNPA1 (ESS Site) | 6289077 | CAGGGC | 83.22 |
| Sironi_motif2 (ESS Site) | 6289086 | CGCGAGG | 61.17 |
| Sironi_motif1 (ESS Site) | 6289086 | CGCGAGGC | 75.30 |
| ESS_hnRNPA1 (ESS Site) | 6289089 | GAGGCT | 77.98 |
| Sironi_motif2 (ESS Site) | 6289091 | GGCTGTG | 60.01 |
| Sironi_motif2 (ESS Site) | 6289094 | TGTGAAG | 66.68 |
| Sironi_motif2 (ESS Site) | 6289113 | TGCTTGG | 63.08 |
| Sironi_motif2 (ESS Site) | 6289114 | GCTTGGC | 62.52 |
| Sironi_motif2 (ESS Site) | 6289117 | TGGCGGA | 61.96 |
| Sironi_motif1 (ESS Site) | 6289123 | ACAGAAGA | 64.77 |
| Sironi_motif2 (ESS Site) | 6289126 | GAAGAGG | 60.55 |
| Sironi_motif1 (ESS Site) | 6289126 | GAAGAGGT | 89.73 |
| ESS_hnRNPA1 (ESS Site) | 6289127 | AAGAGG | 70.84 |
| Sironi_motif2 (ESS Site) | 6289128 | AGAGGTG | 73.70 |
| ESS_hnRNPA1 (ESS Site) | 6289129 | GAGGTG | 74.41 |
| Sironi_motif2 (ESS Site) | 6289130 | AGGTGGG | 73.16 |
| Sironi_motif2 (ESS Site) | 6289131 | GGTGGGT | 85.35 |
| Sironi_motif2 (ESS Site) | 6289139 | TGTGTGA | 72.75 |
| Sironi_motif1 (ESS Site) | 6289141 | TGTGAGGC | 65.65 |
| Sironi_motif2 (ESS Site) | 6289141 | TGTGAGG | 83.58 |
| Sironi_motif2 (ESS Site) | 6289143 | TGAGGCT | 68.47 |
| ESS_hnRNPA1 (ESS Site) | 6289144 | GAGGCT | 77.98 |
| ESS_hnRNPA1 (ESS Site) | 6289150 | TAGAAC | 78.70 |
| Sironi_motif2 (ESS Site) | 6289160 | TCTGGAG | 66.68 |
| Sironi_motif1 (ESS Site) | 6289161 | CTGGAGGG | 76.30 |
| Sironi_motif2 (ESS Site) | 6289162 | TGGAGGG | 72.80 |

| | | | |
|---|---|---|---|
| Sironi_motif2 (ESS Site) | 6289163 | GGAGGGT | 82.30 |
| ESS_hnRNPA1 (ESS Site) | 6289164 | GAGGGT | 87.98 |
| Sironi_motif2 (ESS Site) | 6289167 | GGTTGAG | 72.55 |
| Sironi_motif2 (ESS Site) | 6289170 | TGAGCAG | 63.63 |
| ESS_hnRNPA1 (ESS Site) | 6289181 | TAATGC | 66.08 |
| Sironi_motif3 (ESS Site) | 6289212 | CTTACCAA | 69.36 |
| ESS_hnRNPA1 (ESS Site) | 6289223 | TAACGC | 66.08 |
| Sironi_motif2 (ESS Site) | 6289227 | GCTGGTG | 63.60 |
| Sironi_motif2 (ESS Site) | 6289230 | GGTGATG | 63.60 |
| Sironi_motif2 (ESS Site) | 6289232 | TGATGCT | 60.51 |
| Sironi_motif2 (ESS Site) | 6289238 | TGTTGGG | 92.53 |
| Sironi_motif2 (ESS Site) | 6289239 | GTTGGGA | 69.67 |
| Sironi_motif2 (ESS Site) | 6289263 | TGCTGGT | 67.93 |
| Sironi_motif2 (ESS Site) | 6289264 | GCTGGTG | 63.60 |
| Sironi_motif1 (ESS Site) | 6289265 | CTGGTGGC | 62.62 |
| Sironi_motif2 (ESS Site) | 6289266 | TGGTGGC | 69.96 |
| Sironi_motif2 (ESS Site) | 6289267 | GGTGGCC | 70.48 |
| Sironi_motif3 (ESS Site) | 6289273 | CTTCTCAT | 62.47 |

**ESM Table 5.** Consensus sequences starting from the ATG of the four alternative splicing variants deriving from the c.316-1G>A-carrying allele. Lowercase letters highlighted in cyan refer to the skipped sequence in the wild type isoform.

| **WFS1 splice variant** | **Consensus sequence** |
|---|---|
| NM_006005.3_c.316del | ATGGACTCCAACACTGCTCCGCTGGGCCCCTCCTGCCCACAGCCCCCGCCAGCACCGCAGCCCCAGGCGCGTTCCCGACTCA<br>ATGCCACAGCCTCGTTGGAGCAGGAGAGGAGCGAAAGGCCCCGAGCACCCGGACCCCAGGCTGGCCCTGGCCCTGGTGTTAG<br>AGACGCAGCGGCCCCCGCTGAACCCCAGGCCCAGCATACCAGGAGCCGGGAAAGAGCAGACGGCACCGGGCCTACAAAGGGA<br>GACATGGAAATCCCCTTTGAAGAAGTCCTGGAGAGGGCCAAGGCCGGGGACCCCAAGGCACAGACTGAGgTGGGGAAGCACT<br>ACCTGCAGTTGGCCGGCGACACGGATGAAGAACTCAACAGCTGCACCGCTGTGGACTGGCTGGTCCTCGCCGCGAAGCAGGG<br>CCGTCGCGAGGCTGTGAAGCTGCTTCGCCGGTGCTTGGCGGACAGAAGAGGCATCACGTCCGAGAACGAACGGGAGGTGAGG<br>CAGCTCTCCTCCGAGACCGACCTGGAGAGGGCCGTGCGCAAGGCAGCCCTGGTCATGTACTGGAAGCTCAACCCCAAGAAGA<br>AGAAGCAGGTGGCCGTGGCGGAGCTGCTGGAGAATGTCGGCCAGGTCAACGAGCACGATGGAGGGGCGCAGCCAGGCCCCGT<br>GCCCAAGTCCCTGCAGAAGCAGAGGCGGATGCTGGAGCGCCTGGTCAGCAGCGAGTCCAAGAACTACATCGCGCTGGATGAC<br>TTTGTGGAGATCACTAAGAAGTACGCCAAGGGCGTCATCCCCAGCAGCCTGTTCCTGCAGGACGACGAAGATGATGACGAGC<br>TGGCGGGGAAGAGCCCTGAGGACCTGCCACTGCGTCTGAAGGTGGTCAAGTACCCCCTGCACGCCATCATGGAGATCAAGGA<br>GTACCTGATTGACATGGCCTCCAGGGCAGGCATGCACTGGCTGTCCACCATCATCCCCACGCACCACATCAACGCGCTCATC<br>TTCTTCTTCATCATCAGCAACCTCACCATCGACTTCTTCGCCTTCTTCATCCCGCTGGTCATCTTCTACCTGTCCTTCATCT<br>CCATGGTGATCTGCACCCTCAAGGTGTTCCAGGACAGCAAGGCCTGGGAGAACTTCCGCACCCTCACCGACCTGCTGCTGCG<br>CTTCGAGCCCAACCTGGATGTGGAGCAGGCCGAGGTTAACTTCGGCTGGAACCACCTGGAGCCCTATGCCCATTTCCTGCTC<br>TCTGTCTTCTTCGTCATCTTCTCCTTCCCCATCGCCAGCAAGGACTGCATCCCCTGCTCGGAGCTGGCTGTCATCACCGGCT<br>TCTTTACCGTGACCAGCTACCTGAGCCTGAGCACCCATGCAGAGCCCTACACGCGCAGGGCCCTGGCCACCGAGGTCACCGC<br>CGGCCTGCTATCGCTGCTGCCCTCCATGCCCTTGAATTGGCCCTACCTGAAGGTCCTTGGCCAGACCTTCATCACCGTGCCT<br>GTCGGCCACCTGGTCGTCCTCAATGTCAGCGTCCCGTGCCTGCTCTATGTCTACCTGCTCTATCTCTTCTTCCGCATGGCAC<br>AGCTGAGGAATTTCAAGGGCACCTACTGCTACCTTGTGCCCTACCTGGTGTGCTTCATGTGGTGTGAGCTCTCCGTGGTCAT<br>CCTGCTGGAGTCCACCGGCCTGGGGCTGCTCCGCGCCTCCATCGGCTACTTCCTCTTCCTCTTTGCCCTCCCCATCCTGGTG<br>GCCGGCCTGGCCCTGGTGGGCGTGCTGCAGTTCGCCCGGTGGTTCACGTCTCTGGAGCTCACCAAGATCGCAGTCACCGTGG<br>CGGTCTGTAGTGTGCCCCTGCTGTTGCACTGGTGGACCAAGGCCAGCTTCTCTGTGGTGGGGATGGTGAAGTCCCTGACGCG<br>GAGCTCCATGGTCAAGCTCATCCTGGTGTGGCTCACGGCCATCGTGCTGTTCTGCTGGTTCTATGTGTACCGCTCAGAGGGC<br>ATGAAGGTCTACAACTCCACACTGACCTGGCAGCAGTATGGTGCGCTGTGCGGGCCACGCGCCTGGAAGGAGACCAACATGG<br>CGCGCACCCAGATCCTCTGCAGCCACCTGGAGGGCCACAGGGTCACGTGGACCGGCCGCTTCAAGTACGTCCGCGTGACTGA<br>CATCGACAACAGCGCCGAGTCTGCCATCAACATGCTCCCGTTCTTCATCGGCGACTGGATGCGCTGCCTCTACGGCGAGGCC<br>TACCCTGCCTGCAGCCCTGGCAACACCTCCACGGCCGAGGAGGAGCTCTGTCGCCTTAAGCTGCTGGCCAAGCACCCCTGCC<br>ACATCAAGAAGTTCGACCGCTACAAGTTTGAGATTACCGTGGGCATGCCATTCAGCAGCGGCGCTGACGGCTCGCGCAGCCG<br>CGAGGAGGACGACGTCACCAAGGACATCGTGCTGCGGGCCAGCAGCGAGTTCAAAAGCGTGCTGCTCAGCCTGCGCCAGGGC<br>AGCCTCATCGAGTTCAGCACCATCCTGGAGGGCCGCCTGGGCAGCAAGTGGCCTGTCTTCGAGCTCAAGGCCATCAGCTGCC<br>TCAACTGCATGGCCCAGCTCTCGCCCACCAGGCGGCACGTGAAGATCGAGCACGACTGGCGCAGCACCGTGCATGGCGCCGT<br>GAAGTTCGCCTTCGACTTCTTTTTCTTCCCATTCCTGTCGGCGGCCTGA |

| | |
|---|---|
| NM_006005.3_c.316_460del | ATGGACTCCAACACTGCTCCGCTGGGCCCCTCCTGCCCACAGCCCCCGCCAGCACCGCAGCCCCAGGCGCGTTCCCGACTCA<br>ATGCCACAGCCTCGTTGGAGCAGGAGAGGAGCGAAAGGCCCCGAGCACCCGGACCCCAGGCTGGCCCTGGCCCTGGTGTTAG<br>AGACGCAGCGGCCCCCGCTGAACCCCAGGCCCAGCATACCAGGAGCCGGGAAAGAGCAGACGGCACCGGGCCTACAAAGGGA<br>GACATGGAAATCCCCTTTGAAGAAGTCCTGGAGAGGGCCAAGGCCGGGGACCCCAAGGCACAGACTGAGgtggggaagcact<br>acctgcagttggccggcgacacggatgaagaactcaacagctgcaccgctgtggactggctggtcctcgccgcgaagcaggg<br>ccgtcgcgaggctgtgaagctgcttcgccggtgcttggcggacagaagagGCATCACGTCCGAGAACGAACGGGAGGTGAGG<br>CAGCTCTCCTCCGAGACCGACCTGGAGAGGGCCGTGCGCAAGGCAGCCCTGGTCATGTACTGGAAGCTCAACCCCAAGAAGA<br>AGAAGCAGGTGGCCGTGGCGGAGCTGCTGGAGAATGTCGGCCAGGTCAACGAGCACGATGGAGGGGCGCAGCCAGGCCCCGT<br>GCCCAAGTCCCTGCAGAAGCAGAGGCGGATGCTGGAGCGCCTGGTCAGCAGCGAGTCCAAGAACTACATCGCGCTGGATGAC<br>TTTGTGGAGATCACTAAGAAGTACGCCAAGGGCGTCATCCCCAGCAGCCTGTTCCTGCAGGACGACGAAGATGATGACGAGC<br>TGGCGGGGAAGAGCCCTGAGGACCTGCCACTGCGTCTGAAGGTGGTCAAGTACCCCCTGCACGCCATCATGGAGATCAAGGA<br>GTACCTGATTGACATGGCCTCCAGGGCAGGCATGCACTGGCTGTCCACCATCATCCCCACGCACCACATCAACGCGCTCATC<br>TTCTTCTTCATCATCAGCAACCTCACCATCGACTTCTTCGCCTTCTTCATCCCGCTGGTCATCTTCTACCTGTCCTTCATCT<br>CCATGGTGATCTGCACCCTCAAGGTGTTCCAGGACAGCAAGGCCTGGGAGAACTTCCGCACCCTCACCGACCTGCTGCTGCG<br>CTTCGAGCCCAACCTGGATGTGGAGCAGGCCGAGGTTAACTTCGGCTGGAACCACCTGGAGCCCTATGCCCATTTCCTGCTC<br>TCTGTCTTCTTCGTCATCTTCTCCTTCCCCATCGCCAGCAAGGACTGCATCCCCTGCTCGGAGCTGGCTGTCATCACCGGCT<br>TCTTTACCGTGACCAGCTACCTGAGCCTGAGCACCCATGCAGAGCCCTACACGCGCAGGGCCCTGGCCACCGAGGTCACCGC<br>CGGCCTGCTATCGCTGCTGCCCTCCATGCCCTTGAATTGGCCCTACCTGAAGGTCCTTGGCCAGACCTTCATCACCGTGCCT<br>GTCGGCCACCTGGTCGTCCTCAATGTCAGCGTCCCGTGCCTGCTCTATGTCTACCTGCTCTATCTCTTCTTCCGCATGGCAC<br>AGCTGAGGAATTTCAAGGGCACCTACTGCTACCTTGTGCCCTACCTGGTGTGCTTCATGTGGTGTGAGCTCTCCGTGGTCAT<br>CCTGCTGGAGTCCACCGGCCTGGGGCTGCTCCGCGCCTCCATCGGCTACTTCCTCTTCCTCTTTGCCCTCCCCATCCTGGTG<br>GCCGGCCTGGCCCTGGTGGGCGTGCTGCAGTTCGCCCGGTGGTTCACGTCTCTGGAGCTCACCAAGATCGCAGTCACCGTGG<br>CGGTCTGTAGTGTGCCCCTGCTGTTGCACTGGTGGACCAAGGCCAGCTTCTCTGTGGTGGGGATGGTGAAGTCCCTGACGCG<br>GAGCTCCATGGTCAAGCTCATCCTGGTGTGGCTCACGGCCATCGTGCTGTTCTGCTGGTTCTATGTGTACCGCTCAGAGGGC<br>ATGAAGGTCTACAACTCCACACTGACCTGGCAGCAGTATGGTGCGCTGTGCGGGCCACGCGCCTGGAAGGAGACCAACATGG<br>CGCGCACCCAGATCCTCTGCAGCCACCTGGAGGGCCACAGGGTCACGTGGACCGGCCGCTTCAAGTACGTCCGCGTGACTGA<br>CATCGACAACAGCGCCGAGTCTGCCATCAACATGCTCCCGTTCTTCATCGGCGACTGGATGCGCTGCCTCTACGGCGAGGCC<br>TACCCTGCCTGCAGCCCTGGCAACACCTCCACGGCCGAGGAGGAGCTCTGTCGCCTTAAGCTGCTGGCCAAGCACCCCTGCC<br>ACATCAAGAAGTTCGACCGCTACAAGTTTGAGATTACCGTGGGCATGCCATTCAGCAGCGGCGCTGACGGCTCGCGCAGCCG<br>CGAGGAGGACGACGTCACCAAGGACATCGTGCTGCGGGCCAGCAGCGAGTTCAAAAGCGTGCTGCTCAGCCTGCGCCAGGGC<br>AGCCTCATCGAGTTCAGCACCATCCTGGAGGGCCGCCTGGGCAGCAAGTGGCCTGTCTTCGAGCTCAAGGCCATCAGCTGCC<br>TCAACTGCATGGCCCAGCTCTCGCCCACCAGGCGGCACGTGAAGATCGAGCACGACTGGCGCAGCACCGTGCATGGCGCCGT<br>GAAGTTCGCCTTCGACTTCTTTTTCTTCCCATTCCTGTCGGCGGCCTGA |
| NM_006005.3_c.316_456del | ATGGACTCCAACACTGCTCCGCTGGGCCCCTCCTGCCCACAGCCCCCGCCAGCACCGCAGCCCCAGGCGCGTTCCCGACTCA<br>ATGCCACAGCCTCGTTGGAGCAGGAGAGGAGCGAAAGGCCCCGAGCACCCGGACCCCAGGCTGGCCCTGGCCCTGGTGTTAG<br>AGACGCAGCGGCCCCCGCTGAACCCCAGGCCCAGCATACCAGGAGCCGGGAAAGAGCAGACGGCACCGGGCCTACAAAGGGA<br>GACATGGAAATCCCCTTTGAAGAAGTCCTGGAGAGGGCCAAGGCCGGGGACCCCAAGGCACAGACTGAGgtggggaagcact<br>acctgcagttggccggcgacacggatgaagaactcaacagctgcaccgctgtggactggctggtcctcgccgcgaagcaggg |

| | |
|---|---|
| | ccgtcgcgaggctgtgaagctgcttcgccggtgcttggcggacagaAGAGGCATCACGTCCGAGAACGAACGGGAGGTGAGG<br>CAGCTCTCCTCCGAGACCGACCTGGAGAGGGCCGTGCGCAAGGCAGCCCTGGTCATGTACTGGAAGCTCAACCCCAAGAAGA<br>AGAAGCAGGTGGCCGTGGCGGAGCTGCTGGAGAATGTCGGCCAGGTCAACGAGCACGATGGAGGGGCGCAGCCAGGCCCCGT<br>GCCCAAGTCCCTGCAGAAGCAGAGGCGGATGCTGGAGCGCCTGGTCAGCAGCGAGTCCAAGAACTACATCGCGCTGGATGAC<br>TTTGTGGAGATCACTAAGAAGTACGCCAAGGGCGTCATCCCCAGCAGCCTGTTCCTGCAGGACGACGAAGATGATGACGAGC<br>TGGCGGGGAAGAGCCCTGAGGACCTGCCACTGCGTCTGAAGGTGGTCAAGTACCCCCTGCACGCCATCATGGAGATCAAGGA<br>GTACCTGATTGACATGGCCTCCAGGGCAGGCATGCACTGGCTGTCCACCATCATCCCCACGCACCACATCAACGCGCTCATC<br>TTCTTCTTCATCATCAGCAACCTCACCATCGACTTCTTCGCCTTCTTCATCCCGCTGGTCATCTTCTACCTGTCCTTCATCT<br>CCATGGTGATCTGCACCCTCAAGGTGTTCCAGGACAGCAAGGCCTGGGAGAACTTCCGCACCCTCACCGACCTGCTGCTGCG<br>CTTCGAGCCCAACCTGGATGTGGAGCAGGCCGAGGTTAACTTCGGCTGGAACCACCTGGAGCCCTATGCCCATTTCCTGCTC<br>TCTGTCTTCTTCGTCATCTTCTCCTTCCCCATCGCCAGCAAGGACTGCATCCCCTGCTCGGAGCTGGCTGTCATCACCGGCT<br>TCTTTACCGTGACCAGCTACCTGAGCCTGAGCACCCATGCAGAGCCCTACACGCGCAGGGCCCTGGCCACCGAGGTCACCGC<br>CGGCCTGCTATCGCTGCTGCCCTCCATGCCCTTGAATTGGCCCTACCTGAAGGTCCTTGGCCAGACCTTCATCACCGTGCCT<br>GTCGGCCACCTGGTCGTCCTCAATGTCAGCGTCCCGTGCCTGCTCTATGTCTACCTGCTCTATCTCTTCTTCCGCATGGCAC<br>AGCTGAGGAATTTCAAGGGCACCTACTGCTACCTTGTGCCCTACCTGGTGTGCTTCATGTGGTGTGAGCTCTCCGTGGTCAT<br>CCTGCTGGAGTCCACCGGCCTGGGGCTGCTCCGCGCCTCCATCGGCTACTTCCTCTTCCTCTTTGCCCTCCCCATCCTGGTG<br>GCCGGCCTGGCCCTGGTGGGCGTGCTGCAGTTCGCCCGGTGGTTCACGTCTCTGGAGCTCACCAAGATCGCAGTCACCGTGG<br>CGGTCTGTAGTGTGCCCCTGCTGTTGCACTGGTGGACCAAGGCCAGCTTCTCTGTGGTGGGGATGGTGAAGTCCCTGACGCG<br>GAGCTCCATGGTCAAGCTCATCCTGGTGTGGCTCACGGCCATCGTGCTGTTCTGCTGGTTCTATGTGTACCGCTCAGAGGGC<br>ATGAAGGTCTACAACTCCACACTGACCTGGCAGCAGTATGGTGCGCTGTGCGGGCCACGCGCCTGGAAGGAGACCAACATGG<br>CGCGCACCCAGATCCTCTGCAGCCACCTGGAGGGCCACAGGGTCACGTGGACCGGCCGCTTCAAGTACGTCCGCGTGACTGA<br>CATCGACAACAGCGCCGAGTCTGCCATCAACATGCTCCCGTTCTTCATCGGCGACTGGATGCGCTGCCTCTACGGCGAGGCC<br>TACCCTGCCTGCAGCCCTGGCAACACCTCCACGGCCGAGGAGGAGCTCTGTCGCCTTAAGCTGCTGGCCAAGCACCCCTGCC<br>ACATCAAGAAGTTCGACCGCTACAAGTTTGAGATTACCGTGGGCATGCCATTCAGCAGCGGCGCTGACGGCTCGCGCAGCCG<br>CGAGGAGGACGACGTCACCAAGGACATCGTGCTGCGGGCCAGCAGCGAGTTCAAAAGCGTGCTGCTCAGCCTGCGCCAGGGC<br>AGCCTCATCGAGTTCAGCACCATCCTGGAGGGCCGCCTGGGCAGCAAGTGGCCTGTCTTCGAGCTCAAGGCCATCAGCTGCC<br>TCAACTGCATGGCCCAGCTCTCGCCCACCAGGCGGCACGTGAAGATCGAGCACGACTGGCGCAGCACCGTGCATGGCGCCGT<br>GAAGTTCGCCTTCGACTTCTTTTTCTTCCCATTCCTGTCGGCGGCCTGA |
| NM_006005.3_c.271_513del | ATGGACTCCAACACTGCTCCGCTGGGCCCCTCCTGCCCACAGCCCCCGCCAGCACCGCAGCCCCAGGCGCGTTCCCGACTCA<br>ATGCCACAGCCTCGTTGGAGCAGGAGAGGAGCGAAAGGCCCCGAGCACCCGGACCCCAGGCTGGCCCTGGCCCTGGTGTTAG<br>AGACGCAGCGGCCCCCGCTGAACCCCAGGCCCAGCATACCAGGAGCCGGGAAAGAGCAGACGGCACCGGGCCTACAAAGGGA<br>GACATGGAAATCCCCTTTGAAGAAgtcctggagagggccaaggccggggaccccaaggcacagactgaggtggggaagcact<br>acctgcagttggccggcgacacggatgaagaactcaacagctgcaccgctgtggactggctggtcctcgccgcgaagcaggg<br>ccgtcgcgaggctgtgaagctgcttcgccggtgcttggcggacagaagaggcatcacgtccgagaacgaacgggaggtgagg<br>cagctctcctccgagaccgacCTGGAGAGGGCCGTGCGCAAGGCAGCCCTGGTCATGTACTGGAAGCTCAACCCCAAGAAGA<br>AGAAGCAGGTGGCCGTGGCGGAGCTGCTGGAGAATGTCGGCCAGGTCAACGAGCACGATGGAGGGGCGCAGCCAGGCCCCGT<br>GCCCAAGTCCCTGCAGAAGCAGAGGCGGATGCTGGAGCGCCTGGTCAGCAGCGAGTCCAAGAACTACATCGCGCTGGATGAC<br>TTTGTGGAGATCACTAAGAAGTACGCCAAGGGCGTCATCCCCAGCAGCCTGTTCCTGCAGGACGACGAAGATGATGACGAGC |

| | |
|---|---|
| | TGGCGGGGAAGAGCCCTGAGGACCTGCCACTGCGTCTGAAGGTGGTCAAGTACCCCCTGCACGCCATCATGGAGATCAAGGA<br>GTACCTGATTGACATGGCCTCCAGGGCAGGCATGCACTGGCTGTCCACCATCATCCCCACGCACCACATCAACGCGCTCATC<br>TTCTTCTTCATCATCAGCAACCTCACCATCGACTTCTTCGCCTTCTTCATCCCGCTGGTCATCTTCTACCTGTCCTTCATCT<br>CCATGGTGATCTGCACCCTCAAGGTGTTCCAGGACAGCAAGGCCTGGGAGAACTTCCGCACCCTCACCGACCTGCTGCTGCG<br>CTTCGAGCCCAACCTGGATGTGGAGCAGGCCGAGGTTAACTTCGGCTGGAACCACCTGGAGCCCTATGCCCATTTCCTGCTC<br>TCTGTCTTCTTCGTCATCTTCTCCTTCCCCATCGCCAGCAAGGACTGCATCCCCTGCTCGGAGCTGGCTGTCATCACCGGCT<br>TCTTTACCGTGACCAGCTACCTGAGCCTGAGCACCCATGCAGAGCCCTACACGCGCAGGGCCCTGGCCACCGAGGTCACCGC<br>CGGCCTGCTATCGCTGCTGCCCTCCATGCCCTTGAATTGGCCCTACCTGAAGGTCCTTGGCCAGACCTTCATCACCGTGCCT<br>GTCGGCCACCTGGTCGTCCTCAATGTCAGCGTCCCGTGCCTGCTCTATGTCTACCTGCTCTATCTCTTCTTCCGCATGGCAC<br>AGCTGAGGAATTTCAAGGGCACCTACTGCTACCTTGTGCCCTACCTGGTGTGCTTCATGTGGTGTGAGCTCTCCGTGGTCAT<br>CCTGCTGGAGTCCACCGGCCTGGGGCTGCTCCGCGCCTCCATCGGCTACTTCCTCTTCCTCTTTGCCCTCCCCATCCTGGTG<br>GCCGGCCTGGCCCTGGTGGGCGTGCTGCAGTTCGCCCGGTGGTTCACGTCTCTGGAGCTCACCAAGATCGCAGTCACCGTGG<br>CGGTCTGTAGTGTGCCCCTGCTGTTGCACTGGTGGACCAAGGCCAGCTTCTCTGTGGTGGGGATGGTGAAGTCCCTGACGCG<br>GAGCTCCATGGTCAAGCTCATCCTGGTGTGGCTCACGGCCATCGTGCTGTTCTGCTGGTTCTATGTGTACCGCTCAGAGGGC<br>ATGAAGGTCTACAACTCCACACTGACCTGGCAGCAGTATGGTGCGCTGTGCGGGCCACGCGCCTGGAAGGAGACCAACATGG<br>CGCGCACCCAGATCCTCTGCAGCCACCTGGAGGGCCACAGGGTCACGTGGACCGGCCGCTTCAAGTACGTCCGCGTGACTGA<br>CATCGACAACAGCGCCGAGTCTGCCATCAACATGCTCCCGTTCTTCATCGGCGACTGGATGCGCTGCCTCTACGGCGAGGCC<br>TACCCTGCCTGCAGCCCTGGCAACACCTCCACGGCCGAGGAGGAGCTCTGTCGCCTTAAGCTGCTGGCCAAGCACCCCTGCC<br>ACATCAAGAAGTTCGACCGCTACAAGTTTGAGATTACCGTGGGCATGCCATTCAGCAGCGGCGCTGACGGCTCGCGCAGCCG<br>CGAGGAGGACGACGTCACCAAGGACATCGTGCTGCGGGCCAGCAGCGAGTTCAAAAGCGTGCTGCTCAGCCTGCGCCAGGGC<br>AGCCTCATCGAGTTCAGCACCATCCTGGAGGGCCGCCTGGGCAGCAAGTGGCCTGTCTTCGAGCTCAAGGCCATCAGCTGCC<br>TCAACTGCATGGCCCAGCTCTCGCCCACCAGGCGGCACGTGAAGATCGAGCACGACTGGCGCAGCACCGTGCATGGCGCCGT<br>GAAGTTCGCCTTCGACTTCTTTTTCTTCCCATTCCTGTCGGCGGCCTGA |

### EMS Fig. 1

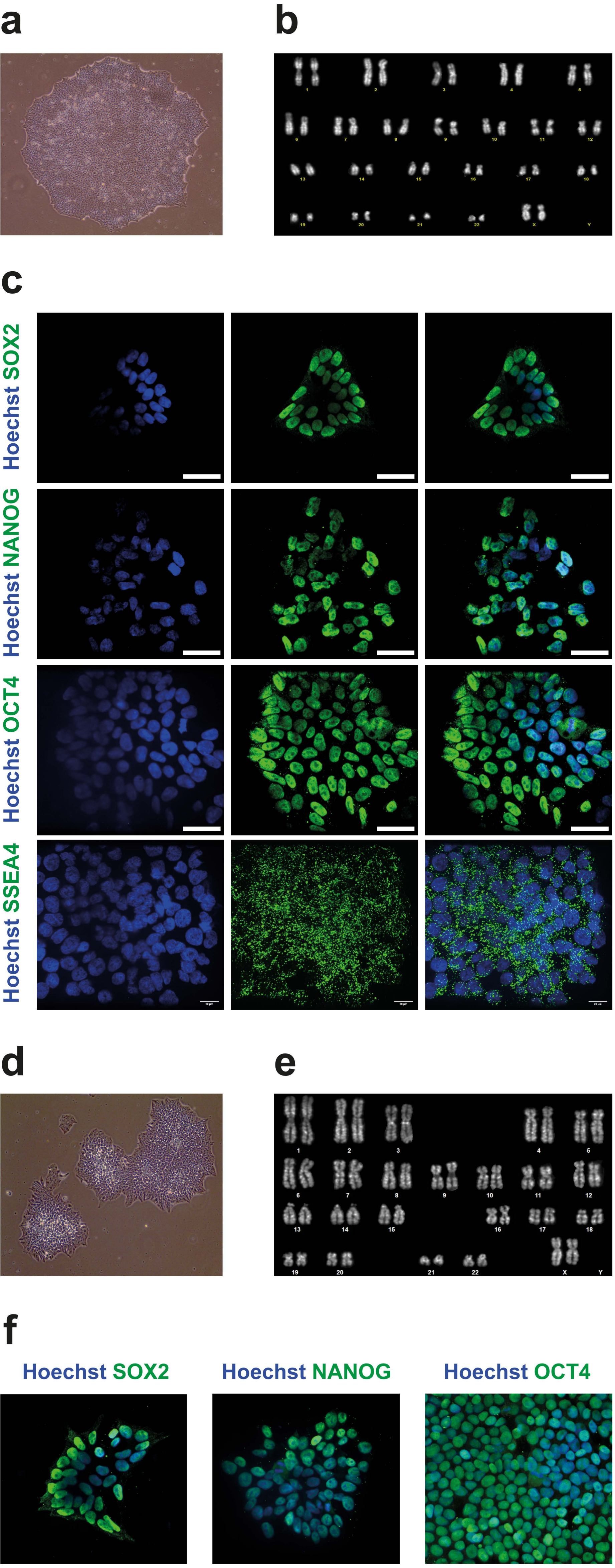


**(a)** Representative image at 10x magnification of WFS1 iPSC colony. **(b)** Representative karyotype of stabilized WFS1 iPSC clones. **(c)** Immunofluorescence analysis of pluripotency markers (SOX2, NANOG, OCT4 and SSEA4) in WFS1 iPSCs. Nuclei were counterstained with Hoechst. Scale bar 40μm. **(d)** Representative image at 10x magnification of WFS1$^{wt/757A>T}$ iPSC colony. **(e)** Representative karyotypes of stabilized WFS1$^{wt/757A>T}$ iPSC clones. **(f)** Immunofluorescence analysis of pluripotency markers (SOX2, NANOG and OCT4) in the WFS1$^{wt/757A>T}$ iPSCs. Nuclei were counterstained with Hoechst. Scale bar 40μm.

### EMS Fig. 2

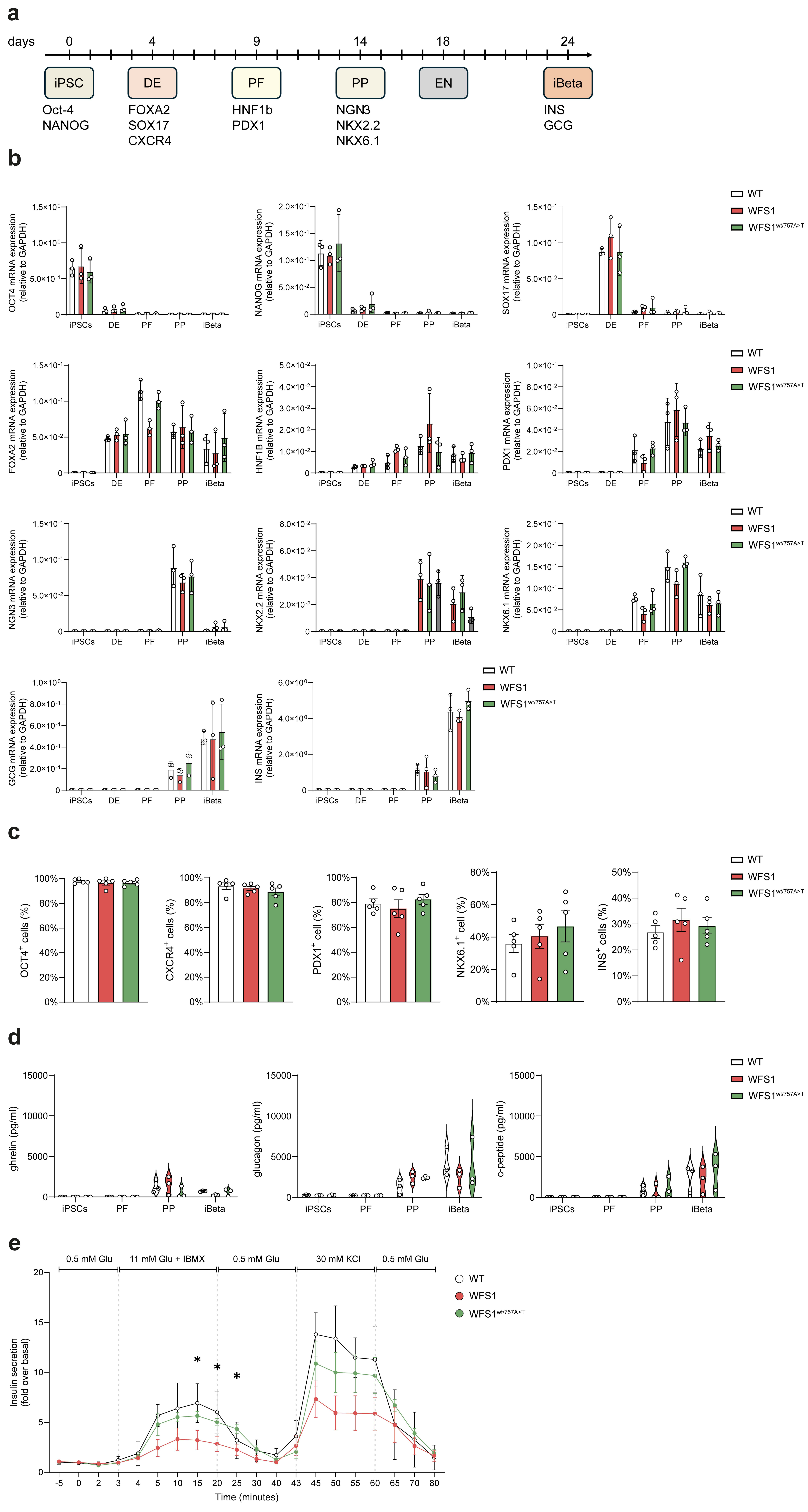


**(a)** Schematic representation of iPSC differentiation process into pancreatic β cells (iBeta), indicating the key markers of each step. DE=Definitive Endoderm; PF=Posterior Foregut; PP=Pancreatic Progenitors; EN=Endocrine Cells. **(b)** Gene expression analysis by TaqMan of markers of pluripotency (OCT4 and NANOG), DE (SOX17 and FOXA2), PF (HNF1B and PDX1), PP (NGN3, NKX2.2 and NKX6.1) and EN/iBeta (GCG and INS) stages of WT, WFS1 and WFS1$^{wt/757A>T}$ iPSCs. Data are plotted as mean±SD. N=3 independent in vitro differentiations. **(c)** FACS analysis of OCT4 in iPSCs, CXCR4 at DE stage, PDX1 at PF stage, NKX6.1 at PP stage, and INS in iBeta expressed as percentage (%) of positive cells. Data are plotted as mean±SD. N=5 independent in vitro differentiations. **(d)** Secretory capacity of iPSC-derived insulin-producing cells. Unstimulated levels of ghrelin, glucagon and C-peptide in the supernatant of WT, WFS1 and WFS1$^{wt/757A>T}$ in iPSCs and at different stages of differentiation (PF, PP and iBeta) were measured by Luminex and expressed as pg/mL. Data are plotted as mean±SD. N=5 independent in vitro differentiations. **(e)** Dynamic perifusion to assess insulin secretion following stimulation with high glucose (11 mM) + IBMX, and 30 mM of KCl. Data are plotted as fold from the baseline, mean±SEM. N=3 (WT) and 5 (WFS1 and WFS1$^{wt/757A>T}$) independent in vitro differentiations. The data were analyzed using two-way ANOVA with Dunn-Šídák correction (*$p<0.05$).

### EMS Fig. 3

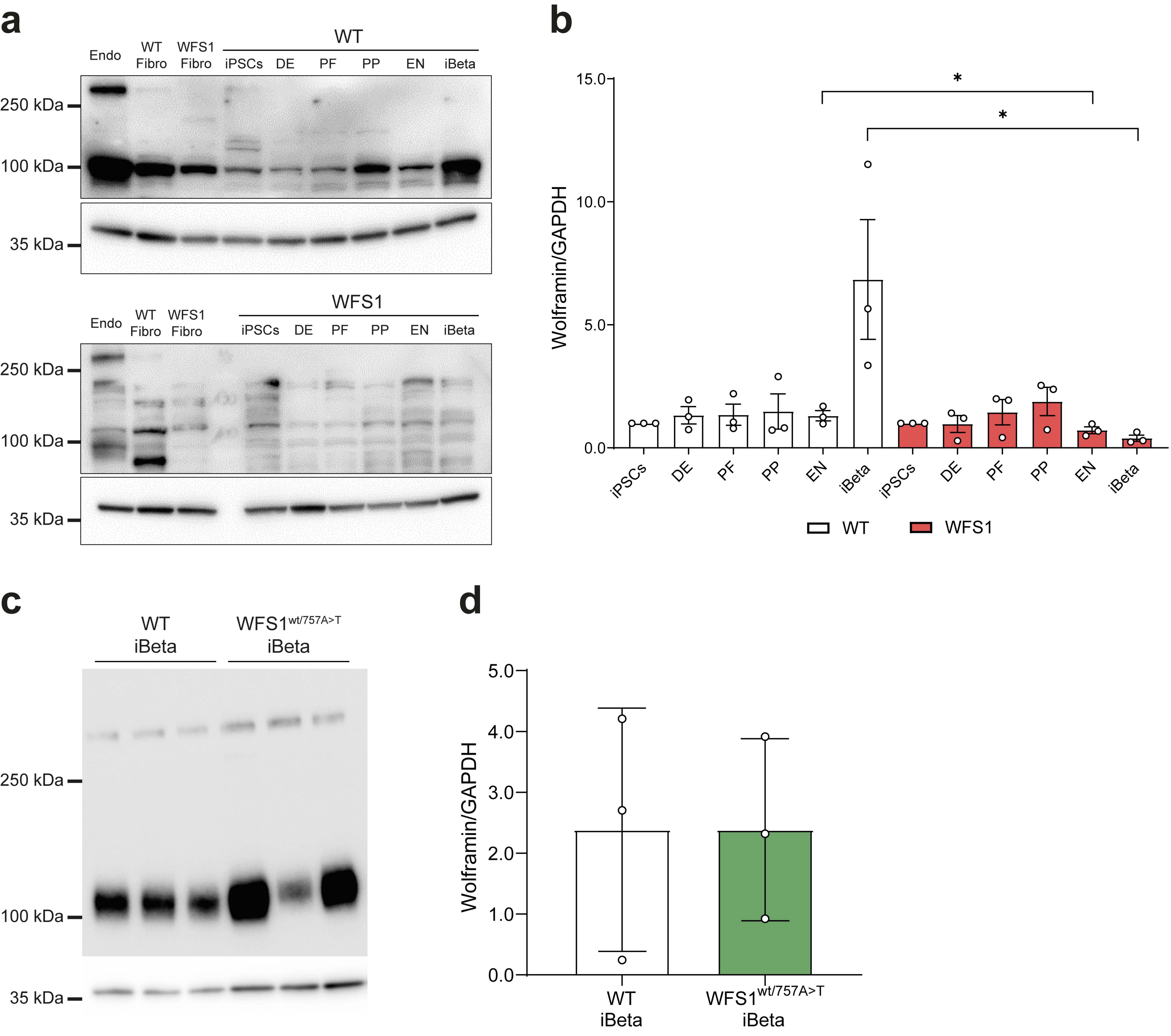


**(a)** Immunoblot analysis of Wolframin expression in WT and WFS1 iPSCs and throughout the differentiation steps (DE, PF, PP, EN, iBeta). EndoC-βH1 cells, WT and WFS1 fibroblasts were used as controls. GAPDH was used as housekeeping gene. **(b)** Relative quantification expressed as Wolframin/GAPDH ratio and expressed as mean±SD. N=3 independent in vitro differentiation. The data were analyzed using the Student's unpaired one-tailed t-test (*p<0.05). **(c)** Immunoblot analysis of Wolframin expression in WT and WFS1$^{wt/757A>T}$ iBeta. GAPDH was used as housekeeping gene. **(d)** Relative quantification expressed as Wolframin/GAPDH ratio and expressed as mean±SD. N=3.

### EMS Fig. 4

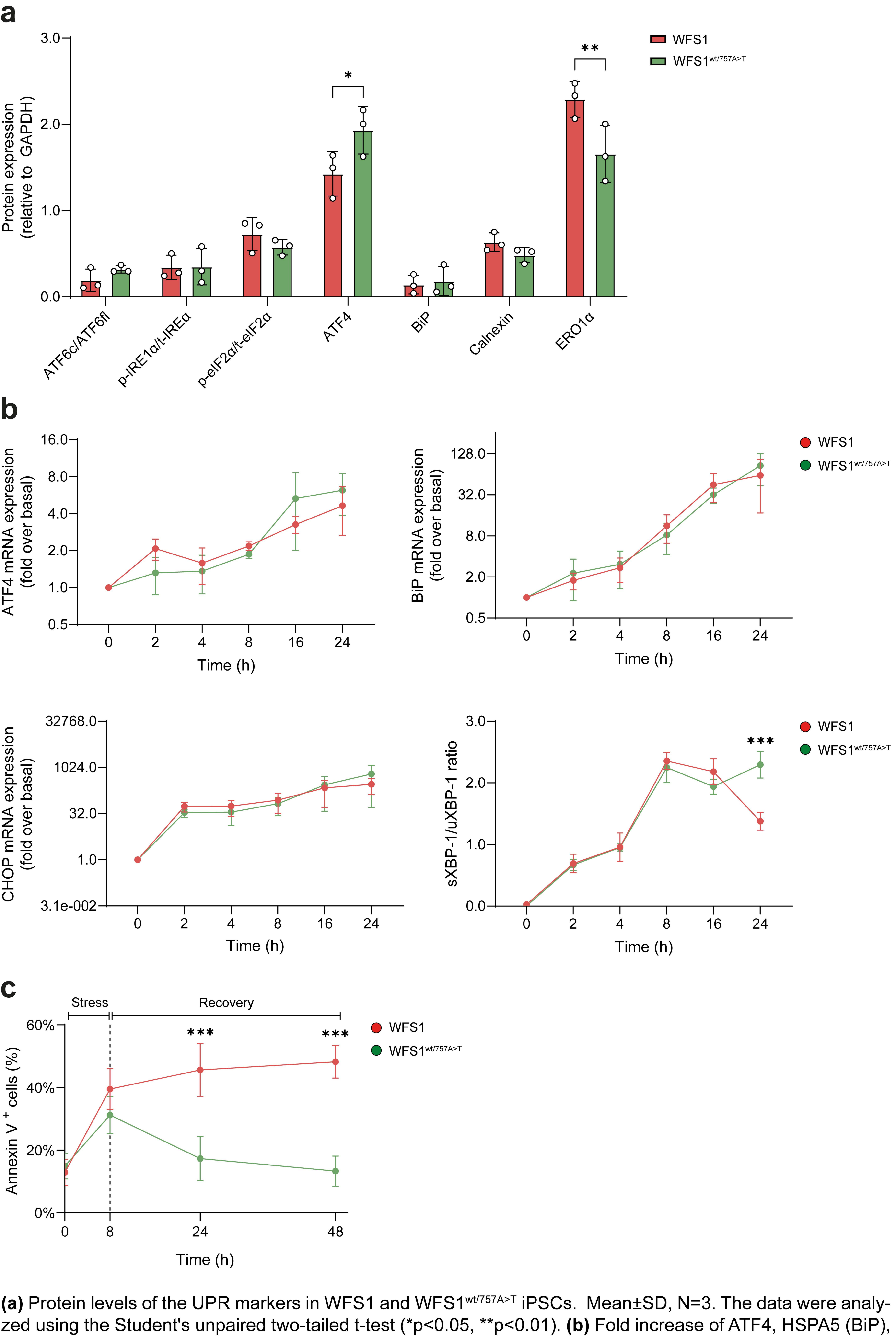


**(a)** Protein levels of the UPR markers in WFS1 and WFS1$^{wt/757A>T}$ iPSCs. Mean±SD, N=3. The data were analyzed using the Student's unpaired two-tailed t-test (*p<0.05, **p<0.01). **(b)** Fold increase of ATF4, HSPA5 (BiP), DDIT3 (CHOP) genes and sXBP1/uXBP1 ratio changes in WFS1 and WFS1$^{wt/757A>T}$ iPSCs after 100 nM TG treatment at the indicated times. Data are plotted as mean±SD. N=3-4 independent experiments. **(c)** Annexin V-positive cell percentages in WFS1 and WFS1$^{wt/757A>T}$ iPSCs during treatment with 100 nM TG and in the recovery period with fresh medium w/o TG. Data are plotted as mean±SEM. N=3. The data were analyzed using two-way ANOVA with Dunn-Šídák correction (***p<0.001).
